# SpinePy enables automated 3D spatiotemporal quantification of multicellular *in vitro* systems

**DOI:** 10.1101/2025.09.10.674634

**Authors:** Ryan G. Savill, Alba Villaronga-Luque, Marc Trani Bustos, Yonit Maroudas-Sacks, Julia Batki, Alexander Meissner, Allyson Q. Ryan, Carl D. Modes, Otger Campàs, Jesse V Veenvliet

## Abstract

Organoids and stem-cell-based embryo models such as gastruloids are powerful systems to quantitatively study morphogenesis and patterning. This requires 3D analysis in reference frames that emerge dynamically over development, but are less stereotypical than *in vivo*. Meaningful statistical comparison and interpretation of biological and physical quantities in space and time — such as signaling activity, gene expression or cell flows — depend on proper quantification in these internal and dynamic coordinate systems, especially in *in vitro* systems that naturally exhibit larger variation. Here, we present a computational framework, packaged as a modular Python toolkit termed SpinePy, that identifies the emergent primary body axis (“spine”) of individual gastruloids and constructs a local, dynamic coordinate system aligned to their morphology. SpinePy enables 3D quantification relative to evolving axes and statistical comparisons of morphodynamics, patterning, and densities across structures with varying geometries. We validate and benchmark SpinePy using both synthetic and experimental gastruloid data, providing practical insights into method performance. Using this framework, we generate 3D patterning maps from gastruloids formed with different initial cell numbers (*N*_0_). This reveals distinct patterning classes that are better explained by gastruloid volumes than by *N*_0_. While demonstrated in gastruloids, SpinePy is broadly applicable to any multicellular system where analysis relative to evolving internal axes is needed, advancing quantitative and comparative spatial biology.

## Introduction

The advent of advanced 3D *in vitro* model systems that recapitulate key aspects of their *in vivo* counterparts, such as organoids and stem-cell-based embryo models, represents a new frontier in developmental and biomedical research as a much-needed complement to animal models and 2D cultures (1–4). The ability to mimic key aspects of tissue architecture and function — in combination with major technical advancements in 3D imaging (5–9) — has paved the path for high-throughput, high-content studies of normal, perturbed, and pathological conditions. This includes genetic and compound screens, disease modeling, and drug toxicity testing.

Among these models, gastruloids have emerged as a particularly useful system to study key developmental processes in mouse and human, such as axial patterning and morphogenesis (10–15). Gastruloids are generated by coaxing mouse or human pluripotent stem cell aggregates to break symmetry and elongate along one axis, resulting in a distinct, reproducible morphology with self-organized patterning reminiscent of the posterior trunk (10, 13–21) (Fig. 1A). The scalable, accessible, and tractable nature of gastruloids offers unique opportunities to provide quantitative insights into mammalian development, including human, at unprecedented resolution and scale.

**Fig. 1.**
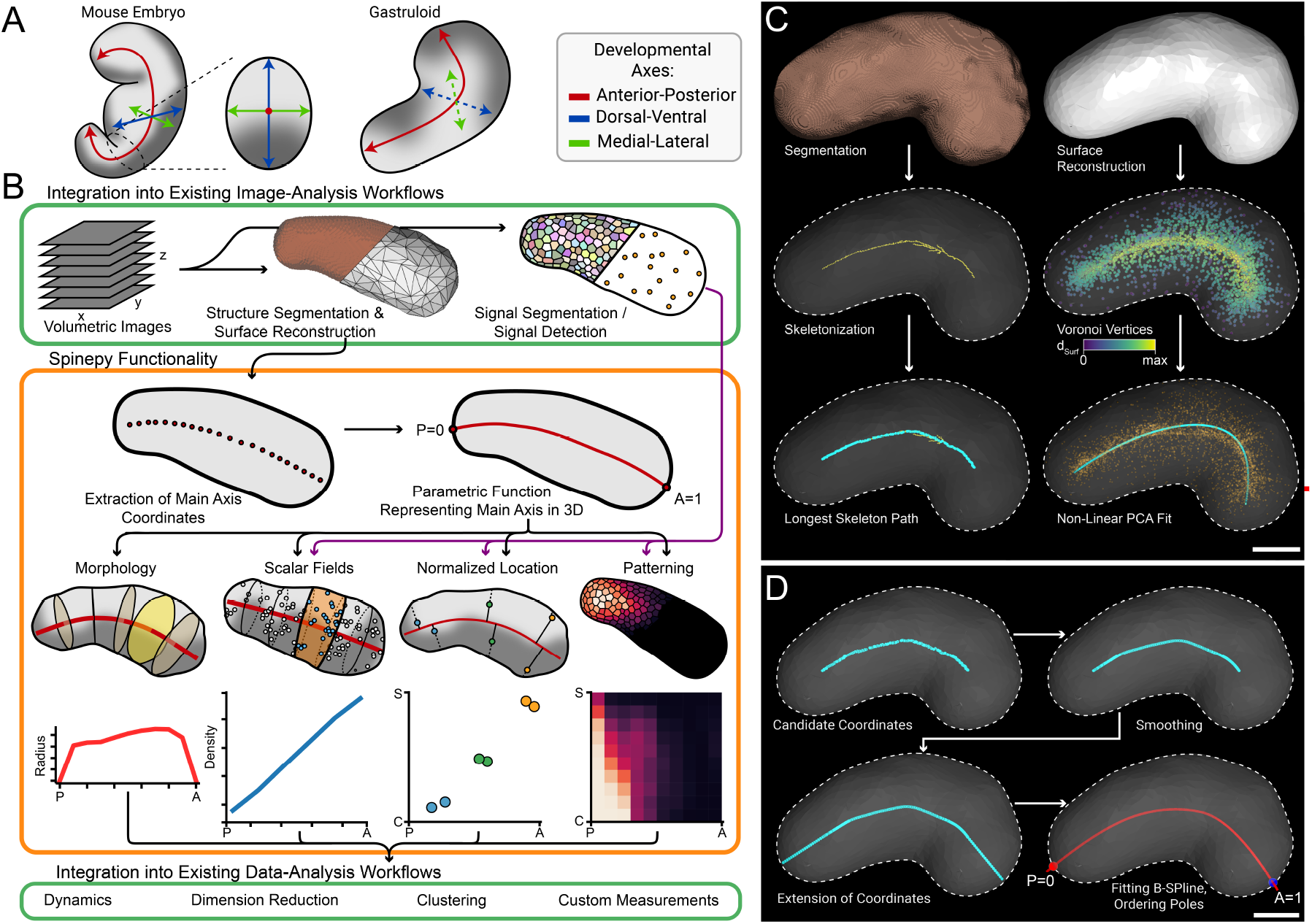
SpinePy Overview and Axis Detection: **A** Schematic of a mouse embryo (left), with transversal section at the trunk level (middle), and of a gastruloid (right). The developmental axes are highlighted. Dashed lines indicate that the capacity to form that axis varies between gastruloids and protocols (4, 16, 19, 34, 35) **B** Overview of the SpinePy functionality. Green boxes highlight external workflow components, orange box highlights SpinePy framework components. **C** 3D visualization of axis detection for a gastruloid at 109 h (see Methods for details, see Fig. S1A for input data). Dashed lines indicate surface mesh boundary, cyan points indicate detected putative spine coordinates Left: skeletonization approach, going from segmented label image (top, brown), to skeletonized image (middle, yellow pixels indicate image skeleton), to the longest path along the image skeleton (cyan). Right: Non-Linear Principal Component Analysis (NLPCA) approach, which takes the surface mesh (top) as input to generate Voronoi vertices colored by their distance to the surface (middle). NLPCA is then fit to the point cloud (bottom, points orange & NLPCA path cyan), yielding a putative spine path. Scale bar: 100 *µ*m. **D** Post-processing of putative spine coordinates. The dashed line indicates the surface reconstruction boundary. Top left: input coordinates generated by the skeletonization approach (cyan). Top right: points after smoothing and resampling using a Savitsky-Golay filtering approach. Bottom left: extension of coordinates to fill the spaces between putative spine endpoints and surface. Bottom right: fit B-spline (red) with indication of poles (P: posterior, A: anterior). Curve was fit after manual ordering of poles so that the spine function of 0 returns the posterior pole and 1 returns the anterior pole. Scale bar: 100 *µ*m.

To fully leverage the potential of organoids and stem-cell-based embryo models, analytical frameworks that allow for statistically robust comparisons across many samples are essential, particularly because these systems exhibit higher intrinsic variation than their *in vivo* counterparts (6, 10, 12, 14, 19, 22–24). This need has sparked the development of image analysis workflows that can deal with the scale and complexity of the data. Wide-field-imaging or maximum-projection-based 2D analysis can be useful to map phenotypic landscapes (22, 25–28), but these approaches result in information loss, can mask intricate patterning, overemphasize surface features, and are in general more error-prone, for example when structures grow out of the plane of imaging. More recently, 3D analysis frameworks have been developed to extract features across multiple scales, from (sub)cellular to whole-organoid levels (5–7, 29). However, these approaches typically operate in the absence of a defined local and dynamic coordinate system, limiting their ability to perform spatial comparisons or compute summary statistics of multiscale features at the whole-organoid level. Without a common reference frame, quantifying and comparing tissue-level patterning in 3D — a critical step for identifying phenotypic variation, interpreting perturbation effects, and uncovering morphodynamic principles across organoid populations — remains challenging.

*In vivo*, such analyses are facilitated by the presence of well-defined and stereotypic (developmental) axes, as well as the stereotypical presence of clear anatomical landmarks (e.g., node, notochord, primitive streak, neural tube), which allow spatial features to be readily interpreted in a common anatomical framework. Accordingly, 3D analysis frameworks for embryos often use these landmarks to align individual embryos to a shared reference frame (30, 31), enabling the identification of expression gradients along developmental axes (32, 33). However, these frameworks are not directly transferable to *in vitro* systems such as gastruloids, which often lack consistent anatomical landmarks and, despite overall reproducibility, exhibit substantial variation in shape and size. This makes it challenging to define a shared reference frame for spatial comparisons.

To address these limitations, we developed SpinePy, a comprehensive 4D (3D+time) imaging analysis framework that automatically detects the main body axis (“spine”) in gastruloids and quantifies multi-scale features — such as morphometry, scalar and vector fields — relative to this axis, directly in the native 3D space. We use synthetic and experimental data to validate and systematically evaluate multiple approaches to extract the spine and quantify signals as a function thereof. This also provides practical insights into methodological performance to guide researchers in selecting the best-suited approach tailored to their data characteristics and analysis needs.

To demonstrate the potential of SpinePy for biological discovery, we extracted 3D patterning maps from gastruloids generated from different initial cell numbers (*N*_0_), followed by dimensionality reduction and unsupervised clustering. This identified distinct gastruloid classes with characteristic marker patterns, which correlate with volumetric growth. Furthermore, benchmarking against synthetic data showed that SpinePy’s 3D analysis robustly distinguishes spatial phenotypes that are missed by conventional 2D maximum projection–based methods, underscoring the importance of quantifying patterning directly in native 3D space.

## Results

### SpinePy Workflow Overview

We describe SpinePy’s general layout, showcasing the capabilities of the framework, its modular architecture, and capacity for seamless integration of common imaging analysis workflows (Fig. 1B). The modular architecture of SpinePy allows flexible incorporation of data at different scales, such as whole structure segmentation, spot detection results, or cellular segmentation, facilitating the integration of SpinePy into existing imaging analysis workflows. SpinePy then uses the gross morphology to detect the “spine” — representing the internal, curvilinear anteroposterior (AP) axis of the gastruloid — and quantifies biological features (using the other input data types) as a function of this coordinate system.

The first step in the SpinePy workflow is to locate the spine — the intrinsic AP axis — of the gastruloid. As gastruloids characteristically elongate along this axis during development (16, 18–21), it can be reliably defined from gross morphology using segmentation labels or surface reconstructions of the entire structure (Fig. 1B, top). SpinePy extracts putative spine coordinates from these inputs, optionally refines them through postprocessing, and fits a smooth B-spline function to the resulting coordinates. This provides a continuous reference along which normalized coordinates (0 at the posterior to 1 at the anterior pole) are assigned. These coordinates enable precise spatial quantification of biological and/or physical features — such as morphology, scalar fields (e.g., density profiles), spatial information, and marker distribution (patterning) maps (Fig. 1B, middle). Subsequently, these high-dimensional feature spaces can be explored through data-driven methods such as dimensionality reduction, clustering, or morphodynamic analyses. Furthermore, users can generate custom measurements tailored to specific biological questions based on the established axis and normalized reference frame (Fig. 1B, bottom).

### Automatic Detection of the Anterior-Posterior Axis in 4D

First, we derive putative spine coordinates using two complementary strategies based on (i) volumetric segmentation and (ii) triangulated surface reconstruction. The segmentation-based approach applies topological skeleton analysis (ToSkA (36)) to a binary gastruloid mask and retrieves the longest geodesic path in the resulting skeleton (Fig. 1C – Left, see Fig. S1A for input data). Points lying within a user-defined margin of the surface can be discarded to refine the path. The surface-based method operates on the vertices of a triangulated mesh. After generating interior seed points via Voronoi tessellation (Methods), we fit a non-linear principal curve (NLPCA) (33, 37, 38) that minimizes the Euclidean distance to the seed set (Fig. 1C, right). Seeds can optionally be weighted or filtered by their normalized surface distance to bias the fit; the resulting path is used as the putative spine. For both strategies, outputs can be manually corrected when automated detection is compromised by incomplete imaging, poor signal-to-noise, or imaging artifacts, which can lead to inaccurate or fragmented reconstructions of the full structure. In such cases users may annotate spine coordinates and proceed with the same downstream workflow.

To refine these putative spine coordinates before fitting a B-spline function to them, post-processing steps are recommended. Firstly, the coordinates have to be spatially ordered, which is achieved via a minimum spanning tree approach. Since skeleton-derived or manual coordinates can show small-scale fluctuations, smoothing can be applied using a Savitzky–Golay filter (Fig. 1 top right, Fig. S1B). To extend the main axis fully toward the posterior and anterior poles of the gastruloid, additional points must be appended from each endpoint toward the surface, guided by the local spine curve tangent at the spine ends (Fig. 1 bottom left). Finally, a B-spline is fitted to these refined coordinates, defining a normalized spatial reference frame (from 0:posterior to 1:anterior) based on intersections with the surface at the gastruloid poles. To ensure correct orientation, anterior or posterior poles can be identified via marker-based staining (e.g., expression of Brachyury (also known as T or TBXT) at the posterior (16, 18–21)) or manual annotation (Fig. 1 bottom right). To ensure consistent pole annotation for time-lapse data, the pole defined at the final time-point is propagated backwards (Methods).

### Validation of Axis Detection Algorithms

To systematically evaluate and validate axis extraction and other analytical components of our SpinePy framework, we generated synthetic gastruloid surface meshes with known curvatures, thickness profiles, and axis coordinates. In short: curved paths with varying degrees of curvature (defined by curve coordinates **c**_*i*_, *i* = 0,…, *n*) were combined with sigmoidal radial profiles **r**_*i*_, *i* = 0,…, *n* to create realistic gastruloid morphologies (Fig. S1C). To simulate realistic variation in the surface mesh, Perlin noise was applied to perturb surface vertices along their normals (Fig. S1D, see Supplemental Note 1 for details).

This provided us with a ground-truth to systematically compare various approaches for extracting the spine: NLPCA applied directly to Voronoi vertices (NLPCA-raw), NLPCA with vertices weighted by distance to the surface (NLPCA-weights), NLPCA with exclusion of vertices exceeding a certain normalized distance threshold (NLPCA-thresh), and skeletonization-based spine detection. To evaluate the performance of each of these methods, we computed the relative errors between the extracted spine and the ground-truth coordinates (Fig. S1E, Methods). Overall, all methods demonstrated good performance in spine extraction. Among them, NLPCA-thresh and skeletonization showed a slight but significant improvement over the other NLPCA approaches (Fig. S1E). Notably, NLPCA-thresh exhibited the greatest consistency, with the lowest fraction of outliers — errors exceeding 1.5 times the 75th percentile — compared to all other methods (Fig. S1E).

To further evaluate the performance of our SpinePy framework, we benchmarked it on real data of fixed gastruloids with more complex morphologies than the synthetic gastruloids, as well as lightsheet live-imaging data. The fixed data covered gastruloids with pronounced curvature and complex morphologies (Fig. S1F). We tasked multiple expert annotators to manually define the AP-axis, and computed the relative error (Methods) for each annotator. This allows us to directly compare algorithmic with manual performance (although we note that the relative error also reflects the error of manual human annotation, so it may capture both algorithmic inaccuracies and human annotation uncertainty). In this context, skeletonization outperformed NLPCA-based methods and performed on-par with the manual annotations (Fig. S1G), whereas NLPCA-based methods performed slightly worse than the manual error baseline. This improved performance likely stems from skeletonization’s more geometrically constrained and unambiguous representation of the spine in thinner, tubular structures, whereas NLPCA-based methods can be challenged by the added complexity in the Voronoi-derived point cloud.

To assess spine extraction accuracy in 4D we used light-sheet live-imaging data. We manually annotated the AP-axis for every fifth time-point for each gastruloid, excluding any time-points with ill-defined axes, yielding a total of 40 annotations (Methods; Fig. S1H). We then compared the spine as automatically defined by SpinePy to the manual annotations as described above. For all spine detection methods, errors were lower or on par with manual-vs-manual errors, with NLPCA-thresh significantly outperforming human annotation (Fig. S1I).

Finally, we evaluated the temporal consistency of automatic axis extraction by measuring the distance between spines at consecutive time-points. This assessment is crucial to ensure that the measured dynamics reflect true biological changes rather than detection variability. NLPCA-thresh significantly outperformed all other methods, and skeletonization exhibited considerable fluctuations — likely due to minor segmentation inconsistencies (Supplemental Video 1). This was reflected in a higher outlier fraction for skeletonization, whereas NLPCA-thresh was the most consistent method with the fewest outliers (Fig. S1J). These results demonstrate the robustness of NLPCA-thresh for tracking continuous temporal changes, highlighting it as the method-of-choice when analyzing time-resolved imaging data.

Collectively, this benchmarking with synthetic and experimental data demonstrates that SpinePy extracts the gastruloid AP axis with human-level — or better — precision, while operating much faster, facilitating automatic, reliable, highthroughput axis annotation in complex 3D and 3D live imaging data.

### Optimization of Orthogonal Slices

To quantify morphology, morphodynamics, or scalar fields along the AP axis, we need to define orthogonal planes evenly spaced along the spine (Fig. 2A - left). The simplest approach would be to define planes, with plane normals corresponding to the local tangent of the spine. However, with such an approach, the orthogonal planes could intersect within the structure for spines with high curvature, leading to overlapping volume segments and erroneous measurements (Fig. 2A - right). To resolve this, we implemented an optimization strategy that adjusts plane normals using a custom loss function. Briefly, intersections of planes within the mesh are detected by casting rays orthogonal to plane normals which are used to detect normalized collision distances (0: intersection at surface; 1: intersection at spine) (Fig. 2B, see Supplemental Note 2 for details). Optimizing rotation of plane normals with this loss function yielded optimized, non-overlapping slicing planes (Fig. 2C). Additionally, the loss function enables filtering of poorly converged optimizations by applying a user-defined threshold.

**Fig. 2.**
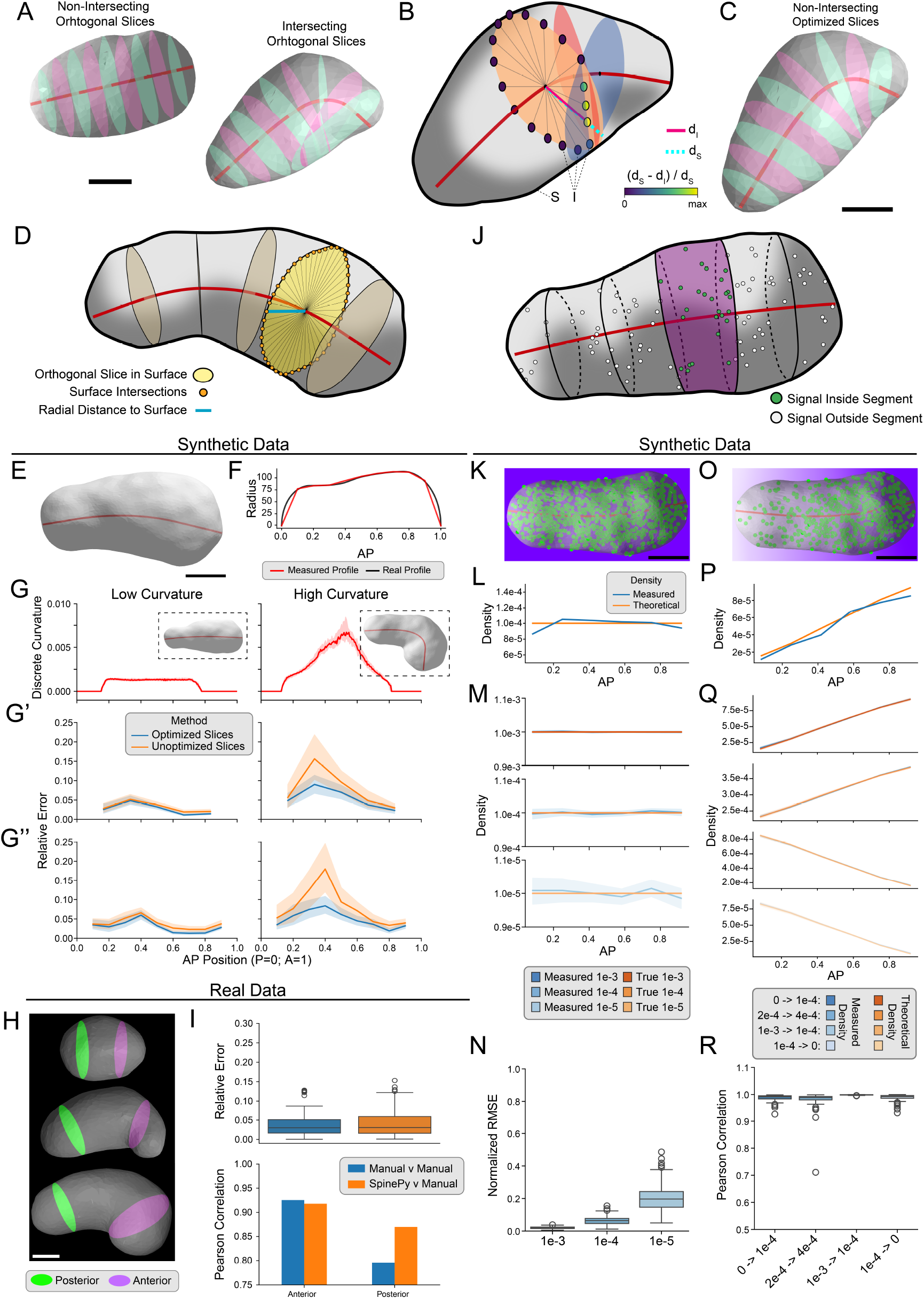
Slicing Optimization & Validation. **A** Visualization of non-intersecting and intersecting orthogonal planes. Spines are shown in red, orthogonal planes are shown in alternating colors (pink and mint) and surface meshes in gray. Scale bar: 100 *µ*m. **B** Visualization of loss determination for a plane of interest. Dark-red: spine; orange: plane of interest; red & blue: intersecting planes; gray: surface mesh (S), intersections (I); magenta: distance to intersection *d*_*I*_ ; dashed-cyan: distance to surface *d*_*S*_ . intersections (I) are color-coded by their intersection loss (see color bar). **C** Structure from A (bottom right) after optimization. **D** Schematic illustrating morphometric measurements. Yellow: plane of interest; blue: radial distance to the surface; orange points: surface intersections. **E** Example of synthetic gastruloid, used to determine profile in F, scale bar: 100 *µ*m. Red: spine; gray: surface mesh. **F** Example profile for the synthetic gastruloid in E. Black: real profile used to generate data; red: profile measured using 9 planes along the spine. **G, G’, G”‘** Curvature and thickness plots along the AP-axis for low (left) and high (right) curvature structures. **G** discrete curvature plot along the AP axis. Insets show representative example structures. Line represents median and shaded area shows 95% confidence interval **G’**,**G”** Relative error for radial thickness plots along the AP-axis using 5 slicing planes (**G’**) or 9 slicing planes **G”**. The relative error is calculated as the absolute error between measured profile and synthetic profile divided by the synthetic profile width. Orange: non-optimized slices; blue: optimized slices. Lines represent mean and shaded regions the 95% confidence interval. **H** Selected manually annotated planes used to measure anterior and posterior thicknesses. Green: posterior plane; purple: anterior plane. Scale bar: 100 *µ*m. **I** Comparison of anterior and posterior radii annotated manually and using SpinePy. Top: Relative error between SpinePy measurement and annotators compared to manual-vs-manual errors. Bottom: Pearson correlation of radii as a function of time. **J** Visualization of the quantification of scalar fields. Red: spine; gray: surface; purple: volume section, green/white: signals inside or outside the volume section, respectively. Signals inside of the segment could represent any biological measurement. **K** Synthetic gastruloid example used to determine the profile in L, scale bar: 100 *µ*m. Dark-red: spine; gray: surface mesh; green: uniform random points; purple: density gradient. **L** Density profile of the synthetic gastruloid shown in K. Blue: simulated density, orange: density measured within each AP section using SpinePy. **M** Uniform density profiles of all synthetic gastruloids. AP: anterior-posterior axis location. Blue: simulated density used; orange: density measured within each AP section using SpinePy. Lines represent mean and shaded regions the 95% confidence interval. **N** Boxplot showing the normalized root mean square error (RMSE) shown for all uniform density samples. Box represents inter quartile range (IQR), whiskers indicate 1.5 * IQR, line denotes median, circles represent outliers. Normalization performed using the simulated density. **O** Synthetic gastruloid example used to determine the profile in P, scale bar: 100 *µ*m. Dark-red: spine; gray: surface mesh; green: random points along density gradient; purple: density gradient. **P** Example gradient density profile of the synthetic gastruloid shown in O. AP: anterior-posterior axis location. Blue: simulated density within AP-slice (see Methods); orange: density measured within each AP section using SpinePy. **Q** Gradient density profiles of all synthetic gastruloids. AP: anterior-posterior axis location. Blue: simulated density within AP-slice (see Methods); orange: density measured within each AP section. Lines represent mean and shaded regions the 95% confidence interval. **R** Boxplot showing Pearson correlations between the simulated density profile and the measured density profile. Box represents inter quartile range (IQR), whiskers indicate 1.5 * IQR, line denotes median, circles represent outliers. Sample sizes: **E, F** - low curvature *n* = 211, high curvature *n* = 80; **I** *n* = 183; **M, N** - *n* = 193; **Q, R** - *n* = 86.

### Morphometric and Morphodynamic Quantifications

To conduct 3D morphology and morphodynamic quantifications along the AP axis, rays are cast at regular intervals around a unit circle from the spine to the surface, perpendicular to the optimized plane normals for all slices (Fig. 2D). Distances along these rays are then recorded, enabling calculation of average radii or diameters per slice in order to generate thickness profiles.

We first validated this measurement method using synthetic data. Since the ground-truth radial profiles used for generation of the synthetic gastruloids are known, we assessed performance by comparing measured profiles against these reference profiles, by computing the relative error (Fig. 2E, F). We compared performances with and without optimization of orthogonal slices for different slice numbers (5 or 9 slices, excluding poles). Since we had observed that optimization of plane normals is more important for higher spine curvatures, we grouped the synthetic gastruloids into two groups with low or high spine curvature (Fig. 2G, Methods).

For structures with lower curvature, both optimized and non-optimized slicing approaches showed similar performances (Fig. 2G’&G”). We note that some degree of error is expected since the reference profiles do not include the noise we add to the surface vertices. For high-curvature structures, the optimized slicing method significantly reduced errors compared to non-optimized slicing (Kruskal-Wallis test *P* = 0.00015), emphasizing the necessity of optimization to obtain accurate measurements. The slice number did not affect errors, although higher slice numbers increased the loss, which can impact the convergence of minimization.

To evaluate method performance on time-lapse data, we used lightsheet live-imaging of gastruloids. Expert annotators were asked to define anterior and posterior planes in 3D to quantify domain-specific radii over time (Fig. 2H). We let SpinePy perform the same task by extracting five optimized slicing planes and averaging the radii from the two anterior and two posterior planes. We then computed relative errors for manual-vs-Spinepy measurements, as well as between annotators (manual-vs-manual) (Fig. 2I; see Methods). Compared to manual annotations, SpinePy achieved similar error levels (average error 0.05) and matched or exceeded manual-vs-manual Pearson correlations.

Altogether, these results show that SpinePy provides robust morphometric and morphodynamic measurements in 3D, with an accuracy on par or better than what can be achieved by manual annotations.

### Volumetric Slicing to Compute Scalar Fields

Within the SpinePy framework, the optimized slicing planes can be used to divide the gastruloid into volumetric sections. Quantifying features such as cell centroids, mitotic events, or marker-positive cells within each slice generates scalar fields along the spine (Fig. 2J), allowing for investigation of the distribution of biological or physical signals in 3D.

To validate scalar field quantification, we analyzed density profiles (*ρ*_*j*_, *j* = 0,…, *n*_sections_) by computing the number of signals per unit of volume in each orthogonal section (Supplemental Video 2). To benchmark against a ground-truth reference, we generated synthetic gastruloids with either uniform density profiles (Fig. 2K-N) or controlled AP density gradients (2O-R) (see Supplemental Note 1 for details). We then used SpinePy to extract the density profile from each synthetic gastruloid and compared these to the ground-truth density. For uniform density profiles, the measured profiles matched the theoretical density profile across three scales of densities (10^*−*3^ to 10^*−*5^, Fig. 2L,M). The normalized root-mean-square error (NRMSE) was lower than 0.05 for denser samples (10^*−*3^) and gradually increased as the density decreased (Fig. 2N). The latter is to be expected, since with lower densities stochastic fluctuations start to dominate the measurement.

Controlled AP gradients with increasing or decreasing density profiles across various density scales were generated by sampling points from an annotated probability field (see Supplemental Note 1 for details; Fig. 2O), and the theoretical density profiles were calculated using the average probability field intensity in each slice (see Supplemental Note 1). For these synthetic AP gradients, we observed a nearly perfect match between the measured and theoretical density profiles (Fig. 2P,Q), with the highest deviations, as expected, occurring at lower densities. This was further confirmed by computing Pearson correlation coefficients between measured and theoretical density, which were >0.98 for all gradient amplitudes and directions, with only few outliers (Fig. 2R).

Altogether, this shows that SpinePy scalar field quantification accurately captures both uniform and gradient density profiles across realistic scales in 3D.

### Tracking Normalized Spatial Information

A key asset of the SpinePy framework is the ability to map spatial locations into a common reference frame, no matter the gastruloids shape, size, or orientation under the microscope. To define this common reference space we can treat a gastruloid as a cylindrical coordinate system (Fig. 3A). In this system each location **l** can be described by a position along the cylinders height *s*, the angle around the cylinder *ϕ* and the position along the cylinders radius *r* (Supplemental Note 3). Since a common reference for the angle *ϕ* at each AP position is not defined, we treat gastruloids as axisymmetric, which allows us to encode position by the two parameters *s* and *r* to retrieve coordinates along the AP-axis and core-to-surface (CS) axis (Fig. 3A’, see Supplemental note 3 for details).

**Fig. 3.**
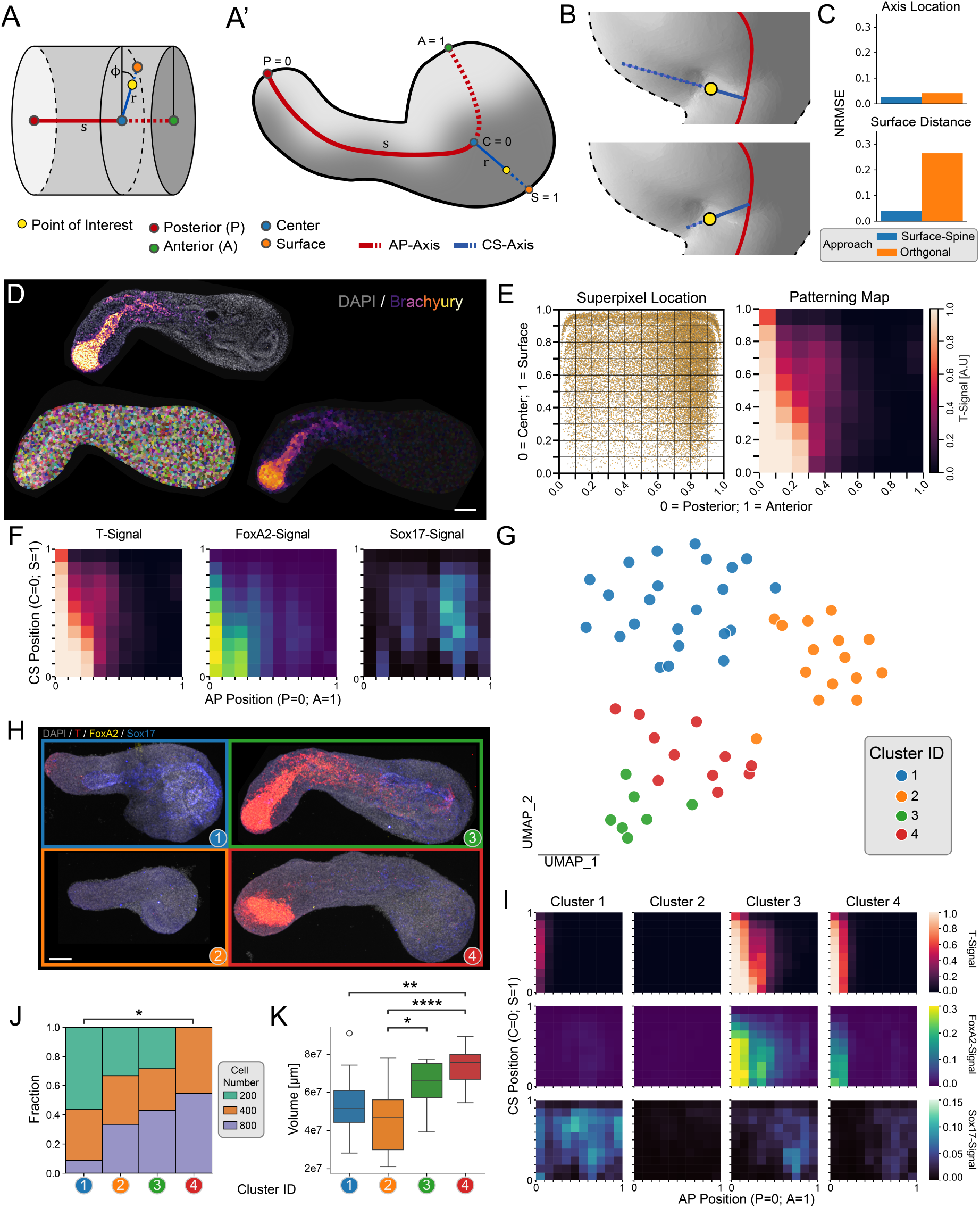
Establishment of a Normalized Reference Frame and Extraction of Patterning Maps: **A, A’** Visualization of normalized spatial quantification. **A** Coordinate system shown for a cylindrical volume. Each point location (yellow) is determined by *c*(*s*) defining the point along the AP axis, *ϕ* the angle of the core-to-surface (CS) axis and *func*(*r*) returning the distance along the CS axis. **A’** The same coordinate system shown for an example gastruloid assuming axisymmetry. The position is now encoded by *c*(*s*) and *func*(*r*) only. **B** Schematic illustration of problematic positions in structures with high curvature. Top: if we determine the position along the AP axis by finding the closest point on the axis (orthogonal approach) the CS-axis position gets skewed, leading to inaccurate measurement of normalized distance. If we instead minimize the distance to both the AP axis and the surface (surface-spine approach) we obtain a slightly different AP-axis position and a more accurate CS-axis position. **C** Error quantification of orthogonal and spine-surface approaches to detect AP and CS distances. Top: normalized root mean squared error (NRMSE) for position along the AP axis when comparing SpinePy-measured locations to manual annotations. Bottom: NRMSE between distance to surface along CS axis and true distance to surface. **D** Gastruloid single *z*-planes sections and quantification of developmental marker signals (here T/Brachyury) using simple linear iterative clustering (SLIC). Top: middle *z*-plane slice of a gastruloid with DAPI signal in gray and T-signal overlaid. Left: superpixel segmentation labels Right: T-signal quantified within the superpixel segmentation labels. Scale bar 100 *µ*m. **E** Left: Super-pixel locations (left) and T patterning map (right) for the example gastruloid shown in D. **F** Patterning maps for all developmental markers (T, FOXA2, SOX17) for the example gastruloid shown in D. **G** Uniform Manifold Approximation and Projection (UMAP) of patterning map features colored by cluster identities. **H** Representative maximum intensity projections of gastruloids for each cluster. Gray: DAPI; red: T; yellow: FOXA2; blue: SOX17. Scale bar 100 *µ*m **I** Average patterning maps for each cluster. **J** Cumulative bar plots showing the fractions of initial cell number conditions (*N*_0_ ) for each cluster. Statistics: Chi-square test, *P* = 0.0157. Post hoc pairwise Chi-square tests were performed with Benjamini-Hochberg FDR correction; * *P* < 0.05 indicates significance. **K** Boxplot showing volumes for each cluster. The box represents the interquartile range (IQR), whiskers indicate 1.5 × IQR, the line denotes the median, and circles represent outliers. Statistics: One-way fixed-effects ANOVA using Type-II sums of squares tested with an F-test, *P* = 1.2 × 10^*−*5^. Post hoc pairwise comparisons were performed using Tukey’s Honest Significant Difference (Tukey-HSD) test. Significance levels: * *P* < 0.05, ** *P* < 0.01, *** *P* < 0.001, **** *P* < 0.0001. Sample sizes: **C** - *n* = 84 Locations in 8 gastruloids; **G, I, J, K** Cluster 1 - *n* = 23, Cluster 2 - *n* = 14, Cluster 3 - *n* = 6, Cluster 4 - *n* = 13.

The simplest strategy to find the AP position is to project the spatial location onto the closest spine point (Supplemental Note 3). Although this “orthogonal” approach is fast and exact for straight or slightly curved segments, it breaks down when the spine bends sharply (Fig. 3B - top). In such regions the nearest spline point is not necessarily the one that best represents the location’s radial path to the surface.

To resolve this ambiguity, we introduced a “spine–surface” optimization. For every candidate spine point we compute two quantities: (i) the Euclidean distance to the spatial location, and (ii) the residual distance from the spatial location to the external surface along the same vector. We then choose the spine point that minimizes the sum of these two distances, effectively balancing proximity to the spine with alignment to the true surface normal (Fig. 3B - bottom, Supplemental Note 3).

To assess performance, we manually annotated the AP coordinate of 84 locations in 8 gastruloids distributed along strongly curved spines (Fig. 3C; see Methods). The spine–surface method reduced AP-position errors slightly relative to the orthogonal approach and, more importantly, decreased CS-axis errors by 50%. We therefore adopted the spine–surface method for all subsequent location-based analyses.

### Quantification of Patterning with Patterning Maps

Motivated by SpinePy’s ability to extract normalized 3D spatial information, we developed a workflow to systematically capture the three-dimensional distribution of developmental markers and generate standardized patterning maps. We first performed 3D segmentation, surface reconstruction, and automatic AP axis detection using the skeletonization approach. Next, we identified labels for signal quantification. The modular design of the SpinePy framework makes it agnostic to the specific choice of labels: it can integrate with advanced deep-learning–based nuclear or cellular segmentation approaches (e.g., StarDist, PlantSeg (39, 40)) when high-resolution imaging is available, or alternatively with superpixel-based methods such as simple linear iterative clustering (SLIC) in cases where imaging does not permit reliable nuclear or cellular segmentation (41).

To facilitate higher throughput, we opted for the latter approach, matching the number of superpixels to the estimated cell count (28) (Methods; (Fig. 3D, top and left)), and used these as labels to quantify intensity features. To minimize artifacts from optical scattering, particularly along the optical (*z*) axis, we analyzed thresholded signals (for example, classified as T^+^ or T^*−*^, Fig. 3D - right; Methods). By detecting normalized positions, each super-pixel can be mapped onto a common reference frame defined by anterior-posterior (AP) and core-to-surface (CS) location (Fig. 3E, left, Supplemental Video 3). Standardized patterning maps were created by aggregating the super-pixel positions into equal bins along each axis and calculating mean signals within bins (Fig. 3E - right). These maps facilitate direct comparisons between samples, addressing the inherent difficulty of deriving summary statistics for samples of variable size and morphology in 3D.

### SpinePy 3D Analysis Outperforms 2D Maximum Projection Based Analysis of Patterning

It is common practice to analyze patterning in gastruloids and organoids after maximum intensity projection (26–28). To benchmark our SpinePy 3D spatial analysis against such 2D approaches, we generated synthetic gastruloids with four distinct posterior T patterning classes: (1) posterior low intensity signal (uniform low), (2) posterior high intensity signal (uniform high), (3) posterior high signal intensity limited to the surface (shell), (4) posterior high signal intensity at the surface and close to the spine (shell+core) (Fig. S2A, Supplemental Videos 4-7, Supplemental Note 1). We then applied both the SpinePy patterning maps workflow and, in parallel, the profile extraction pipeline described based on 2D maximum projections described in (28) (Methods) (Fig. S2A - bottom). Comparison of the Uniform Manifold Approximation and Projection (UMAP) embeddings of the extracted patterning features (Fig. S2B) revealed that the maximum projection based method failed to separate the uniform high, shell and shell+core patterns patterning classes. By contrast, SpinePy 3D analysis correctly resolved all four patterning classes. This was further confirmed by various cluster separation indices calculated in PCA space (Fig. S2C, Methods), as well as investigation of the normalized patterning spaces (Fig. S2C&E). Thus, SpinePy 3D can reveal differences in patterning that are obscured using current state-of-the-art 2D maximum projection based approaches, highlighting the need to quantify patterning in its native 3D space.

### 3D Patterning Maps Reveal Distinct Patterning Classes in Experimental Gastruloids

Previous work has examined the effects of initial cell number (*N*_0_) on gastruloid composition, morphogenesis, and patterning (20, 28, 42, 43). However, none of these studies addressed how *N*_0_ influences patterning in 3D. Encouraged by the improved resolution of SpinePy 3D patterning maps, we therefore investigated an experimental datasets of 56 gastruloids generated from distinct *N*_0_ (200, 400, or 800 mESCs; Methods). We performed immunofluorescent staining of three key developmental transcription factors in 120 h gastruloids: T, FOXA2, and SOX17. In the developing trunk, T is expressed in neuromesodermal progenitors, nascent mesoderm, mesendoderm, and node/notochord (44–46), FOXA2 is expressed in mesendoderm and node/notochord (46–48), and SOX17 in definitive endoderm (49). The combination of these markers thus distinguishes key axial and germ layer derivatives — (neuro)mesodermal progenitors (T^+^/FOXA2^*−*^/SOX17^*−*^), mesendoderm and axial mesoderm (T^+^/FOXA2^+^/SOX17^*−*^), and definitive endoderm (T^*−*^/FOXA2^+*/−*^/SOX17^+^) — providing a spatial overview of tissue organization and patterning within the gastruloid. We then used the superpixel-based workflow to extract 3D patterning maps for individual gastruloids (Fig. 3F).

To explore the patterning maps without imposing prior assumptions, we applied UMAP for dimensionality reduction, and performed Leiden clustering on the first 10 principal components obtained with principal component analysis (PCA) (Fig. S3A; Methods). Projecting scores of the first principal component (PC-0) onto the UMAP embedding suggested patterning heterogeneity within the initial Cluster 3 (Fig. S3B), which we confirmed by visual inspection (Supplementary Video 8), prompting sub-clustering (Methods). This revealed the presence of four distinct gastruloid patterning classes (Fig. 3G, H), underscored by computing cluster-specific mean patterning maps (Fig. 3I). Cluster 1 shows widespread SOX17 signal with an anterior enrichment of FOXA2&SOX17 co-localization, consistent with endoderm specification. Cluster 2 is largely negative for any of the tested markers. Cluster 3 is characterized by a strong T and FOXA2 signal, with extensive T&FOXA2 co-localization at the posterior extending into the anterior core, and anterior SOX17 expression. Similar to Cluster 3, Cluster 4 shows abundant T and FOXA2 signal that is, however, more restricted to the posterior core, and low anterior SOX17 signal (Fig. 3I; Fig. S3D). Biologically, these clusters may reflect distinct stages of axial midline development. Cluster 4, with T and FOXA2 restricted to the posterior core and low anterior SOX17, could resemble an earlier state enriched in node and posterior axial mesoderm progenitors. In contrast, Cluster 3 — with strong T and FOXA2 co-localization extending anteriorly alongside anterior SOX17 expression — is consistent with a later stage where the notochordal plate elongates anteriorly, positioned adjacent to, but distinct from, the emerging foregut endoderm.

One of the strengths of identifying gastruloid classes via patterning maps is that the resulting classes can be further interrogated using measured features that were not part of the clustering process. This can provide insights into gastruloid characteristics correlated with the emergence of these distinct phenotypes. Such analysis revealed that the Cluster 4 phenotype (T&FOXA2 co-localization restricted to posterior core) could only be derived from *N*_0_ = 400 and *N*_0_ = 800 structures (Fig. 3J). In contrast, all other phenotypes were observed for any *N*_0_ (Fig. 3J). Strikingly, for the highly distinct Cluster 2 (low expression of all tested markers) and Cluster 3 (posterior T&FOXA2 co-localization extending into anterior core), the proportion of samples derived from each *N*_0_ condition (200, 400, or 800) was highly similar, indicating no strong dependence on initial cell number (Fig. 3J). However, what separated these phenotypic classes was their volume: Cluster 2 gastruloids were significantly smaller than Cluster 3 gastruloids (Fig. 3K). Similarly, Cluster 4 structures were significantly larger than Cluster 1/2 structures (Fig. 3K). To quantify these relationships, we used pairwise logistic regression and likelihood ratio tests to compare the explanatory power of *N*_0_ and volume in distinguishing cluster identities (Table 1; see Methods). Together, these results show that while some gastruloid phenotypes are linked to initial cell number, others emerge independently of *N*_0_ and instead correlate more strongly with the final gastruloid volume.

**Table 1.** Comparison of logistic regression models across pairwise cluster contrasts evaluating the contributions of standardized volume and initial cell number (*N*_0_ ). Akaike Information Criterion (AIC) values and likelihood ratio test *P* -values (adjusted for multiple testing using the Holm method) indicate the significance of adding each predictor to the models. Significant *P* -values (*P adj*) highlight predictors that improve model fit for distinguishing cluster identities. **** *P* < 0.0001, *** *P* < 0.001.

| cluster_comparison | AIC_volume_only | AIC_N0_only | AIC_full_model | P_N0 | P_volume | Padj_N0 | Padj_volume |
| --- | --- | --- | --- | --- | --- | --- | --- |
| 2 vs 1 | 52.23971 | 52.96933 | 30.89062 | 3.13e-06 | 9.25e-07 | 1.88e-05 (****) | 5.55e-06 (****) |
| 3 vs 1 | 33.79512 | 34.51840 | 36.42307 | 5.04e-01 | 7.58e-01 | 7.07e-01 | 1.00e+00 |
| 4 vs 1 | 33.69457 | 32.32060 | 32.83384 | 8.80e-02 | 2.23e-01 | 2.64e-01 | 6.68e-01 |
| 3 vs 2 | 25.69598 | 33.33656 | 21.33054 | 1.53e-02 | 1.82e-04 | 7.63e-02 | 7.29e-04 (***) |
| 4 vs 2 | 23.21626 | 35.02115 | 20.13335 | 2.90e-02 | 3.97e-05 | 1.16e-01 | 1.98e-04 (***) |
| 4 vs 3 | 25.62565 | 25.83303 | 27.54547 | 3.53e-01 | 5.92e-01 | 7.07e-01 | 1.00e+00 |

We complemented the UMAP analysis with an in-depth analysis of the PCA space which offers a more interpretable lens into the feature space compared to the UMAP analysis. Firstly, we assessed the robustness of clusters corresponding to gastruloid patterning classes. We evaluated cluster separation in PCA space by bootstrapping 100 random subsets of 90% of the data and for each subset, we compared cluster separation indices between non-shuffled and shuffled labels using the first 10 PCs (Fig. S3C; Methods). This analysis demonstrated consistently better separation for the non-shuffled labels, confirming the robustness of the cluster structure. Investigating the first two principal components revealed that the phenotypic classes separated along PC-0 or PC-1, but rarely both simultaneously (Fig. S3E). Overlaying *N*_0_ and volumes revealed that PC-0 primarily captured differences in volume, with higher scores corresponding to larger structures, independent of *N*_0_. In contrast, PC-1 separated structures by *N*_0_, with higher scores corresponding to *N*_0_ = 200 (Fig. S3F,G). This was further supported by correlation analyses (Fig. S3H), which also showed that PC-3 correlated strongly with both *N*_0_ and volume (Fig. S3H), suggesting it captures a third, growth-related dimension (Fig. S3I-K). As the explained variance dropped sharply beyond the fourth component (Fig. S3L), we did not analyze additional PCs.

Further interpreting PCA, we visualized feature loadings as patterning maps (Fig. S3M,N). PC-0 predominantly represents global intensities and co-localization of T and FOXA2 (Fig. S3M,N). In contrast, PC-1 loadings highlight general SOX17 abundance and anterior expression of FOXA2&SOX17 (Fig. S3M,N). PC-3 shows characteristics of Clusters 3 and 4, with high T and FOXA2 colocalization at the posterior and SOX17 mainly confined to the anterior (Fig. S3N,O). Thus, visualizing PCA feature loadings as patterning maps enables spatial interpretation of principal components, linking abstract variance dimensions to biologically meaningful expression domains.

Together, these results highlight the strength of SpinePy’s patterning maps in condensing complex 3D marker expression into concise, quantitative descriptors that can distinguish phenotypic classes and reveal key features influencing gastruloid development.

## Discussion

Our approach addresses a key limitation in the analysis of 3D *in vitro* models such as organoids and stem-cell-based embryo models, which have emerged as powerful tools for studying development and disease (1–4). These systems offer unique advantages in mimicking tissue architecture and enabling high-throughput screening, but present analytical challenges due to phenotypic variation and the lack of stereotypic anatomical features. While 2D projections have been widely used to extract phenotypic information (22, 25–28), they can obscure 3D patterning features (as we show in Fig. S2). Recent 3D frameworks have made progress in extracting multi-scale features (5–7, 29), but typically operate without a common reference frame, limiting their ability to quantify and compare spatial patterns across samples. By introducing SpinePy, we address this gap by enabling the definition of local, dynamic coordinate systems aligned to gastruloid morphology, which can be mapped across samples, allowing robust comparison of spatial locations in complex multicellular structures of varying size and shape. Unlike embryo-focused frameworks that rely on stereotypic anatomical landmarks (30–33), SpinePy requires no prior axis information. Instead, it leverages 3D skeletonization or nonlinear principal component analysis (NLPCA) to extract the curvilinear main body axis from volumetric imaging data and constructs a local, dynamic coordinate system aligned to gastruloid morphology, thereby enabling statistically robust multi-scale quantification of spatial features across samples and conditions.

Our extensive validations using both synthetic and real gastruloids demonstrate that both the 3D skeletonization and the NLPCA approach achieve expert-level or higher accuracy while greatly increasing processing speeds. Moreover, the validations offer detailed insights into how method performance relates to key sample features and data characteristics, such as 3D versus 4D data and gastruloid morphological complexity, providing practical guidance on the optimal choice of approach. Skeletonization excels at capturing the main axis of gastruloids with more complex shapes, whereas NLPCA performs better for time-lapse applications, where maintaining consistency of the reference frame across time points is essential. Scalar field analysis using volume slicing further enables robust quantification of density profiles by providing more consistent regions along the AP axis, while location-based measurements — though more susceptible to local fluctuations — yield higher spatial resolution. Together, these complementary approaches provide flexible options depending on the sample complexity and experimental design.

Implemented as a modular Python package, SpinePy integrates easily into existing image processing workflows and supports high-throughput, high-content analysis of 3D and 4D imaging datasets. For example, it can be readily combined with pipelines, such as Tapenade (7), that extract local cellular features — such as gene expression or nuclear shape. By providing a local, dynamic coordinate system that can be mapped across samples, SpinePy enables these local features to be spatially contextualized. Cell-level properties can then be linked to global tissue organization, thereby advancing our understanding of self-organization in complex multicellular systems like gastruloids. This capability is also valuable for spatial transcriptomics data (17), where SpinePy can serve to map high-dimensional molecular profiles back into a biologically meaningful 3D frame of reference across replicates and conditions. This enables comparative analysis of gene expression domains and their relationship to emergent tissue-scale organization.

Here, we used SpinePy to quantify scalar fields, as in the case of patterning maps. However, our approach can be readily extended to vector fields, allowing the analysis of features such as cell orientation direction, division angles, alignment of cytoskeletal structures or extracellular matrix (ECM) fibers, and even cell flows relative to the body axes. Further extension to tensor fields could enable quantification of, for example, cell shape anisotropy or strain rates.

In its current version, SpinePy focuses on automated identification of the major (anteroposterior) body axis. An extension of SpinePy could incorporate additional biologically relevant axes, such as the dorsal-ventral and medial-lateral axis. A current limitation of SpinePy is its inability to handle structures with multiple axes (22, 43, 50). One possible approach is to model the main axis as a graph with multiple nodes corresponding to anterior and/or posterior branches, assigning each branch its own normalized anterior-posterior length up to the branching point. However, resolving conflicts and ambiguities in the central region where these branches meet remains a difficult problem that requires further methodological development. Another valuable future extension will be full 4D analysis beyond morphodynamics — tracking scalar and vector fields over time — for which SpinePy’s capability to automatically and accurately extract the evolving body axis provides a strong foundation.

Using SpinePy in combination with dimensionality reduction and unbiased clustering, we performed a comprehensive 3D patterning analysis in gastruloids generated from distinct initial cell numbers (*N*_0_). While previous work showed that absolute size (in terms of volume) is directly linked to *N*_0_ and that key developmental marker expression (including T) scales with system size (28), our 3D data demonstrate that even with tightly controlled *N*_0_, gastruloid volumes vary substantially, which associates with distinct patterning classes. This suggests that — although scaling may occur under specific conditions — gastruloid growth and patterning can diverge markedly despite identical *N*_0_. The discrepancy is unlikely to arise solely from limitations of 2D analysis obscuring some *N*_0_-dependent differences (e.g., shell versus shell+core patterns; Fig. S2). Instead, it is likely to reflect the influence of mESC genetic background, mESC maintenance protocols, and/or gastruloid culture conditions, all known to affect developmental outcomes (4, 10, 51–54). Our framework offers a powerful computational approach for systematic, quantitative investigation of how modulation of such system parameters influences (the emergence of) patterning and morphodynamics.

In conclusion, by enabling comprehensive spatial mapping within a unified reference frame, SpinePy provides a basis for the exploration of complex biological and physical spaces. Here, we focused on its applications in gastruloid research. However, SpinePy’s flexible framework can be readily adapted to other stem-cell-based embryo model systems and organoids exhibiting evolving internal axes, such as intestinal and brain organoids (6, 55, 56), providing a much-needed generalizable framework for spatial analysis in complex 3D structures. By facilitating quantitative comparative studies of complex spatial patterning and morphogenesis, SpinePy opens up new possibilities to chart multi-scale organoid morphospaces (12, 23, 57, 58) across normal, perturbed, and pathological states — capturing how spatial patterning and morphogenesis dynamically unfold in different systems, and in health and disease.

## ACKNOWLEDGEMENTS

We thank all members of the Veenvliet lab, members of the SUMO consortium (in particular Nikolaj Gadegaard and Daniel Reumann), Johannes R. Soltwedel and John D. Treado for helpful discussions. We thank Alexandra Schauer, Deniz Conkar and Simone Gasparato for assistance in method validation. We thank the Organoid & Stem Cell and Light Microscopy Facility (MPI-CBG) for assistance. This work was supported by the Max Planck Gesellschaft, a European Innovation Council (EIC) Pathfinder grant under the Horizon Europe Research and Innovation Program (Horizon-EIC-2021-PathfinderChallenges-01 101071203, SUMO), and the Deutsche Forschungsgemeinschaft (DFG, German Research Foundation) under Germany’s Excellence Strategy – EXC 2068 – 390729961– Cluster of Excellence Physics of Life of TU Dresden. Y.M.-S. is supported by the Physics of Life Excellence Postdoctoral Fellows Program from the Cluster of Excellence Physics of Life of TU Dresden. M.T.B. is supported by a Boehringer Ingelheim Fonds PhD Fellowship. This manuscript was prepared using the HenriquesLab bioRxiv Overleaf template, available at https://www.overleaf.com/latex/templates/henriqueslab-biorxiv-template/nyprsybwffws.

## AUTHOR CONTRIBUTIONS

Conceptualization, R.G.S., J.V.V.; data curation, R.G.S.; formal analysis, R.G.S.; funding acquisition, J.V.V.; investigation, A.V.-L., M.T.B., Y.M-S.; methodology, R.G.S.; project administration, J.V.V.; software, R.G.S., A.R., C.D.M.; resources, J.B., A.M., A.Q.R., C.D.M., O.C., J.V.V.; supervision, O.C., J.V.V.; visualization, R.G.S.; writing – original draft, R.G.S. and J.V.V., with input from C.D.M. and O.C.; and writing – review and editing, R.G.S., M.T.B., A.V.-L., Y.M-S., J.B., A.Q.R..

## Materials and Methods

### Experimental Procedures

#### Mouse Embryonic Stem Cell Culture

Mouse Embryonic Stem Cells (mESCs; F1G4 (59)) or NLS-GFP derived thereof (see below) were cultured under Serum+LIF+feeder conditions at 37 ^*°*^C and 5% CO_2_. Cells were maintained in 6 cm plates (Sarstedt, # 83.3901.300) or 6-well plates (Nunc™, TFS 140675) coated with 0.1% gelatin and a layer of mitotically inactive fibroblasts. The culture medium consisted of: 400 mL Knockout Dulbecco’s Modified Eagle’s Medium (DMEM) (4500 mg*/*mL glucose, w/o sodium pyruvate) (Gibco) supplemented with 75 mL ES cell tested FCS, 5 mL 100x L-glutamine (200 nM) (Lonza #BE17-605E), 5 mL 100x penicillin (5000 U*/*mL)/ streptomycin (5000 µg*/*mL), 5 mL 100x non-essential amino acids (Gibco #11140-35), 1 mL 500x *β*-mercaptoethanol (5 µM, 1000x Invitrogen), 5 mL 100x nucleosides (Chemicon) and 5 µL 10000x of Murine Leukemia Inhibitory Factor (LIF) (Sigma-Aldrich, #ESG1107). Medium was refreshed every 24 hours. Cells were passaged every 48 hours using Trypsin-EDTA (Thermofisher, #11560626) in a dilution of 1:10. Cells were always passaged twice prior to gastruloid generation. Regular tests were conducted to ensure mESCs were mycoplasma-negative, by PCR of the supernatant medium after culturing a well of mESCs for three days without refreshing medium.

#### Generation of NLS-GFP Reporter mESCs

Wild-type F1G4 cells were used to generate the NLS-GFP reporter mESCs. The targeting vector contained the CAG promoter, SV40 NLS, EGFP coding sequence and SV40 polyadenylation signal, flanked by sequences homologous to the Rosa26 locus (genomic coordinates (mm10) chr6:113,076,061-113,078,081 and chr6:113,075,226-113,076,060 for left and right homology arm respectively). This was co-transfected with a plasmid encoding Cas9 (PX459; Addgene, 62988) and containing a gRNA targeting the Rosa26 locus (gRNA sequence: ACTGGAGTTGCAGATCACGA) using FuGENE HD transfection reagent (Promega, E2311) as per the manufacturer’s instructions. Briefly, 400,000 mESCs were plated the day before transfection. For the transfection, 8 µg of each plasmid DNA was diluted in 125 µl Opti-MEM (Thermo Fisher Scientific, 31985062); 25 µl of FuGENE reagent (room temperature) was diluted with 100 µl Opti-MEM. The diluted FuGENE was added to the diluted DNA, incubated at room temperature for 15 min and added dropwise to the cells. The medium was changed the next day. GFP-expressing mESCs were isolated by FACS 48 h after transfection and single-cell-derived clones stably expressing GFP were picked and expanded (NLS-GFP mESCs).

#### Generation of Mouse Gastruloids

Mouse gastruloids were generated as described previously (16, 60). Briefly, feeder-free mESCs were prepared by sequential plating on 0.1% gelatin-coated 6-well plates and incubated at 37 ^*°*^C and 5% CO_2_ for 25, 20, and 15 minutes consecutively. Cells were washed with 5 mL pre-warmed PBS supplemented with MgCl_2_ and CaCl_2_ (PBS++), centrifuged at 300 g for 5 minutes, and then washed again with 5 mL pre-incubated NDiff227 medium (Takara). After pelleting at 300 g for 5 minutes and gentle trituration in 500 µL NDiff227, cells were counted using an automated cell counter (Invitrogen™Countess™3 FL Automated Cell Counter) and diluted in NDiff227. Subsequently, 30 µL of this suspension (corresponding to *N*_0_ = 200, 400 or 800 cells) were seeded into each well of ultra-low attachment U-bottom 96-well plates (Corning #7007) and incubated at 37 ^*°*^C and 5% CO_2_. After 48 h, aggregates were treated with 3 µM CHIR99021 diluted in 150 µL NDiff227. At 72 h and 96 h, 150 µL of NDiff227 medium was removed and replaced with 150 µL of pre-incubated NDiff227. For gastruloids generated from NLS-GFP+ mESCs for lightsheet live imaging, the same protocol was followed with the following exception: 35 µL of the cell suspension was plated per well, corresponding to *N*_0_ = 300.

#### Whole-mount Immunofluorescence

Samples were collected using a wide-bore P200 pipette tip and transferred to an 8-well glass-bottom plate (Ibidi #80827). They were rinsed three times with PBS++/BSA (PBS containing MgCl_2_ and CaCl_2_ [Sigma-Aldrich #D8662-6X500ML] and supplemented with 0.5% BSA), followed by three washes with PBS. Fixation was carried out with 4% PFA for 1 hour at 4 ^*°*^C, after which samples were washed three more times in PBS. Permeabilization was performed using PBS-T (PBS with 0.05% (v/v) Triton X-100), followed by overnight blocking in a solution of PBS with 0.05% (v/v) Triton X-100 and 10% fetal calf serum (FCS). Gastruloids were incubated with primary antibodies (1:250 dilution) for 72–96 hours. Subsequently, samples were washed three times with PBS-T and incubated overnight in blocking solution. The following day, secondary antibodies (1:500 dilution) were added and samples were incubated for 24 hours. Afterward, gastruloids were washed three times with blocking solution, once with PBS-T, and then post-fixed in 4% PFA for 20 minutes to 1 hour. Samples were then rinsed twice in 0.02 M Phosphate Buffer (PB) (25 mM NaH2PO4 + 75 mM Na2HPO4, pH 7.4) and embedded in low-melting point agarose to stabilize them for imaging. Finally, samples were cleared overnight using RIMS (133% (w/v) Histodenz (Sigma-Aldrich, #D2138) diluted in 0.02 M Phosphate Buffer supplemented with 0.01% Tween). A list of all the antibodies used in this manuscript can be found in Table 2 and 3.

**Table 2.**
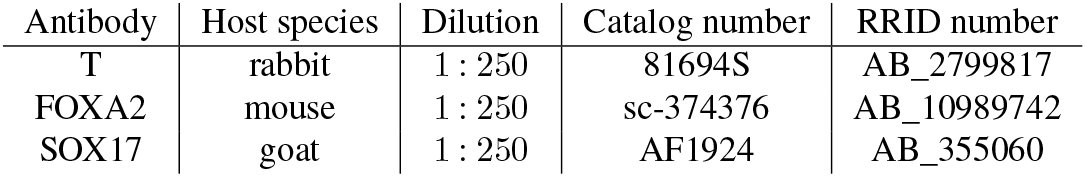
Primary antibodies

| Antibody | Host species | Dilution | Catalog number | RRID number |
| --- | --- | --- | --- | --- |
| T | rabbit | 1 : 250 | 81694S | AB_2799817 |
| FOXA2 | mouse | 1 : 250 | sc-374376 | AB_10989742 |
| SOX17 | goat | 1 : 250 | AF1924 | AB_355060 |

**Table 3.** Secondary antibodies and dyes

| Antibody | Wavelength | Dilution | Catalog number | RRID Number |
| --- | --- | --- | --- | --- |
| DAPI | 405 | 1 : 20,000 | 10236276001 | – |
| Donkey anti-mouse | Alexa 647+ | 1 : 500 | A32787 | AB_2762830 |
| Donkey anti-goat | Alexa 546 | 1 : 500 | 2306813 | AB_2534103 |
| Donkey anti-rabbit | Alexa 488 | 1 : 500 | A21206 | AB_2535792 |

#### Imaging of Fixed Gastruloids

After clearing, samples were imaged on a Spinning Disc microscope (Olympus IXplore SpinSR using Yokogawa W1 with Borealis laser source, iXon Ultra 888 camera, Olympus 10x/0.4 U Plan SApo Air objective and appropriated laser/filters for 405 nm, 488 nm, 568 nm and 640 nm emission fluorophores). Images were acquired with a *z*-step of 9 µm and a pixel size of 0.65 µm^2^.

#### Lightsheet Live Imaging of Mouse Gastruloids

Live imaging of mouse gastruloid axial elongation for 14 hours (96 h - 110 h after aggregation) was performed using a Viventis LS1 lightsheet microscope with an incubation chamber pre-set at 37 ^*°*^C and 5% CO_2_. Viventis FEP Cuvettes were washed with 70% EtOH, rinsed with water, and washed with 70% EtOH again, and then exposed to UV for 30 min under a cell culture hood, in order to sterilize them. Under a tissue culture hood, the FEP cuvettes were then rinsed with 1 ml of anti-adherence rinsing solution (STEMCELL™ Technologies, 07010) for at least 30 min, and subsequently washed three times with NDiff227 medium. Cuvettes with Ndiff227 medium were then placed in an incubator at 37 ^*°*^C and 5% CO_2_ for pre-incubation. 94 h gastruloids that had started to break symmetry (as evident by a tear drop-like shape (60)) were selected for imaging, and transferred from 96-well plates onto the sterilized and pre-incubated Viventis cuvettes using p200 wide-bore pipette tips. Up to 8 different gastruloids could be transferred into a Viventis cuvette and live imaged for every experiment. Subsequently, gastruloids were manually positioned in the cuvettes under a stereoscope in a tissue culture hood, using p200 wide-bore pipette tips. Imaging was performed with a Nikon Apo 25x (1.1 NA) water immersion objective, using an Andor Zyla sCMOS (VSC-12371) camera set at full range, and acquiring 2048x2048 pixels/frame, with 1 µm *z*-steps, imaging every 10 min for 14 h. A 488 nm laser line was used for excitation of GFP, using an OD2 filter and a GFP 525/50 BP filter.

### Computational Procedures

#### Image Preprocessing and Surface Reconstruction

Images from both fixed and live samples were first rescaled to an isotropic voxel size of 2 µm. For **fixed samples**, to correct for the refractive index mismatch between the air objective and the clearing solution (RIMS), the *z*-height was adjusted by a scale factor of 1.48. The nuclear DAPI signal was normalized by rescaling intensity values between 0 and 1 using the 0.1 and 99.9 percentiles as minimum and maximum, with clipping of values outside this range. Gastruloid boundaries were sparsely annotated to train a random-forest-based pixel classifier for segmentation. The largest connected label was retained, and holes were closed by binary closing: dilating with a spherical structuring element (radius = 10 µm) followed by erosion (radius = 14 µm). Remaining holes were filled (61). For **live samples**, after rescaling, nuclear signals were smoothed using a Gaussian blur (kernel size 8) to remove noise, then segmented using Otsu’s thresholding method (62). Only the largest connected component was retained as the segmentation mask. Surface reconstruction for both sample types involved adding a border of zero-valued pixels around the segmentation mask to avoid edge artifacts. The label image was converted into a surface mesh using the marching cubes algorithm. The resulting mesh was softly smoothed using a windowed Sinc kernel. Mesh decimation differed slightly: fixed sample meshes were reduced by a factor of 20, whereas live sample meshes were decimated to 1% of their original vertices. Both meshes then underwent further smoothing via a moving least squares approach, with radius parameters of 20 µm for fixed and 60 µm for live samples. Finally, the initial border offset was subtracted from the mesh vertices.

#### Spine Extraction - Skeletonization Approach

To extract putative spine coordinates using the skeletonization approach we first take the whole structure segmentation and perform skeletonization as implemented in scikit-image (63) before extracting the longest skeleton path using functions implemented in ToSkA (36). The points along this path can then be used for further processing.

#### Spine Extraction - Non-Linear Principal Component Analysis Approach

For non-linear principal component analysis (NLPCA) based spine extraction first a point-cloud is generated to fit the NLPCA. To extract a point-cloud representing the core, the Voronoi diagram of the surface vertices is derived. We first performed Delaunay meshing to turn surface vertices into a set of tetrahedra used to efficiently calculate the Voronoi diagram. Using the surface mesh we then filtered out any vertices outside of the mesh, and calculate the normalized distances 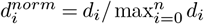 of the Voronoi-vertices to the mesh. This allowed us to either weigh or filter the Voronoi-vertices by a normalized distance threshold, in order to restrain the point-cloud to the core. This point-cloud is then used to fit a NLPCA, returning an optimized path through the point-cloud used as putative spine coordinates.

#### Ordering of Coordinates for Spine Detection

To use the extracted putative coordinates for spine detection we first ensured correct ordering. To achieve this without the need of end-coordinates, we took a graph-based approach in which we generated a fully connected weighted graph with all coordinates as nodes and all distances between coordinates as edge weights. From this graph a minimum spanning tree was generated, and all Dijkstra-distances between leaf nodes were computed. The final path was then defined as the path between the nodes with the highest Dijkstra-distance.

#### Smoothing

Some coordinates have sharp transitions between points, especially in case of discrete methods for extracting coordinates, such as skeletonization or upon manual correction of putative spine coordinates. To smooth these coordinates we use Savitsky-Golay filtering (implemented in SciPy (61)). These filters require equally spaced coordinates, which we achieved by linear interpolation between points prior to smoothing. The degree of smoothing is controlled by the window size and can be adjusted depending on the dataset.

#### Spine Extension

Since the putative spine coordinates do not extend to the surface mesh, we added additional coordinates to fill the space between the putative path and the surface with additional coordinates. These are placed along a vector **v**:

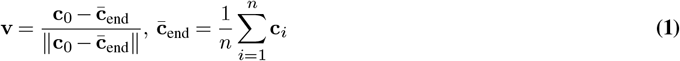

with **c**_0_ defining the coordinate at each end of the spine end, and 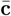 the average of *n* end coordinates. The intersection with the surface mesh is found for the path along this vector and a variable number of points are evenly placed between the spine and surface along the path, either defined by a set number of points or by a density value.

#### Re-parameterizing B-splines

After fitting a parametric B-spline to the data, we re-parameterized it to ensure that evenly spaced parameter values yield evenly spaced points in 3D space. Since direct re-parameterization of B-splines is non-trivial, we implemented a translation function that maps uniform values in [0, 1] to parameter values for the B-spline that define evenly spaced coordinates along the spine between the two poles. This was achieved by sampling parameters between the two poles along the spine with high density and creating an interpolating function for the densely sampled points.

#### Pole Detection

To ensure that axis parameters of 0 and 1 always correspond to the posterior and anterior poles, respectively, the coordinates used for B-spline fitting can be re-ordered either automatically or manually. Automatic detection is based on the quantification of developmental marker gene profiles with known polarity (such as T) along the spine. The pole with higher marker gene signal levels can then be defined as the posterior or anterior pole (depending on the marker gene). Manual annotation of poles is also possible within Napari (REF). For time-lapse data, we employed a semi-automatic method: the posterior pole was annotated for the last time-point *t*_*n*_, when morphology is indicative of the AP polarity. This annotation is then backtracked throughout the time-lapse — for each frame *t*_*i*_ the posterior pole detected at *t*_*i*+1_ is used as a reference to order the coordinates.

#### Manual Spine Annotation for Validation

Three experts in gastruloid biology were tasked to generate a set of points representing the main axis of gastruloids using the DAPI (fixed) or NLS-GFP (live) channel of volumetric image data. Using the 3D image viewer Napari users annotated points and refined their annotations until these matched the perceived spine. To ensure continuity of the annotated axis points, they were ordered using the minimum spanning tree method described above.

#### Validation of AP Axis Detection

For the generation of synthetic gastruloid data we picked *N* = 50 parameters *s*_*i*_ = *i/N* (*i* = 0 … *N* ) on the predicted axis **c**_pred_ to obtain evenly spaced points **p**_*i*_ = **c**_pred_(*s*_*i*_) along the predicted axis. For each point **p**_*i*_ we determined the nearest point **g**_*i*_ on the ground-truth axis and defined the distance to the ground-truth as:

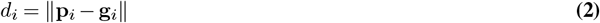

We then calculated the normalized error as *δ*_*i*_ = *d*_*i*_*/r*_*i*_, with *r*_*i*_ representing the radius we used to generate synthetic gastruloids; we report the median of 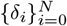.

For the dataset comprising fixed gastruloids with complex morphologies as well as the time-lapse dataset, we used annotations generated by three experts annotators as the ground-truth axis **g**_*i*_ and calculated *d*_*i*_ as in Eq. 2. Since in this case we do not have a ground-truth profile to normalize the distances, we casted the ray **g**_*i*_ + *t* (**p**_*i*_*−* **g**_*i*_) to the surface and let **q**_*S*_ be the first hit. The normalized distance is then defined as:

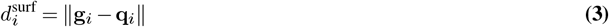

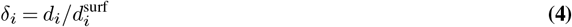

We report the median of 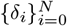 and repeat the measurements for each annotator. All manual annotators were compared to each other to enable a comparison between the manual-vs-manual error and the error of the spine extraction methods.

#### Slice Optimization

The details for the determination of the total loss ℒ are reported in Supplemental Note 2. We used the total loss to first assess whether optimization is needed (i.e., ℒ> 0) and then optimized slice rotation using the Broy-den–Fletcher–Goldfarb–Shanno algorithm in SciPy (61). When the optimization did not converge (i.e., ℒ> threshold) we repeated the optimization procedure or discarded the sample.

For the final loss, we set a threshold of ℒ/*n*_planes_ *<* 5, above which the optimization was deemed unsuccessful and the resulting sample or time point discarded from the analysis.

#### Morphometric and Morphodynamic Measurements

To measure the morphology for a plane parametrized by an origin **p**_*j*_ and normal **n**_*j*_ we casted *n*_ray_ rays in a circle along the plane and measured the average distance to the surface as:

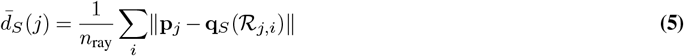

with **q**_*S*_ representing the surface intersection along rays ℛ(*j, i*) casted from the spine in a unit circle, perpendicular to the plane normal **n**_*j*_. See Supplemental Note 2 for details.

#### Discrete Curvature Measurements

To quantify curvature in synthetic gastruloids we measured discrete curvatures using the scaled and resampled coordinates **c**_*i*_, *i* = 0,…, *n* representing the spines used to generate the synthetic data. The discrete curvature *κ*_*j*_ between each coordinate was calculated as:

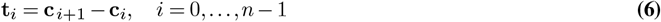

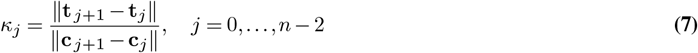

To classify structures into high- and low-curvature groups, we computed the average curvature along the entire axis for each structure and assigned them based on whether this value fell below or above the 50th percentile of the distribution.

#### Validation of Morphometric Measurements with Synthetic Data

For the synthetic gastruloids, we had access to the distance profiles *r*_*i*_ that were used to generate the data. To evaluate the accuracy of the measured radial profiles, we compared the measured radius 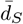 at the AP position to the corresponding synthetic profile. For each slice, we determined the spine parameter *s* corresponding to the synthetic and measured profile and quantified the relative error (RE) as:

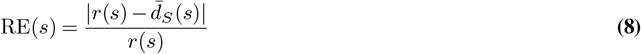

We then analyzed this error along the spine.

#### Validation of Morphometric and Morphodynamic Measurements with Manual Annotations

We asked two experts to manually define planes at the anterior and posterior ends by selecting axis positions and adjusting plane orientations. For each annotated plane, we computed a centroid based on the surface intersections and calculated an average radius as the mean distance from the centroid to these intersections. To determine the anterior and posterior radii using SpinePy we measured morphology as in Eq. 5 using five slices and defined the anterior and posterior radii as the average of the first two and last two slices, respectively. We then computed the relative error between manually annotated and measured radii, as well as inter-annotator error, using the relative error metric defined in Eq. 8.

#### Generations of Synthetic Density Data and Validation of Scalar Field Measurements

Supplemental Note 1 describes the synthetic density data generation in detail. Density along the AP axis *ρ*_*k*_, *k* = 0,…, *n*_sec_ (with *n*_sec_ representing the number of volume sections) was quantified by counting signals within each segment and dividing by the total volume of the corresponding segment. We estimated the error for each section by computing the normalized root mean square error (NRMSE):

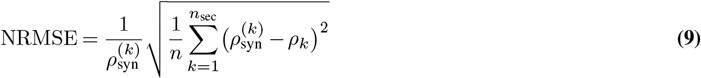

with 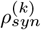 representing the theoretical density level within the slice *k*. For uniform density data this corresponds to the total density used to generate points, and for gradient density evaluation this corresponds to the average density within each slice (see Supplemental Note 2 for details).

#### Validation of Normalized Position

To evaluate the spatial information we defined 8-12 locations along the AP axis of a subset of fixed gastruloids stained with the nuclear marker DAPI. Two experts then tasked to adjust a slider (in Napari) that determined the corresponding AP position, while reviewing the vectors between the AP-position and spatial location. We then computed the normalized root mean squared error between the predicted axis position *ŝ* _*i*_ and the mean annotated axis position *s*_*i*_ as:

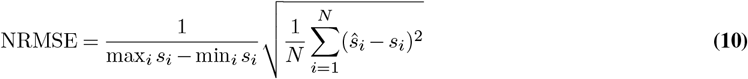

For the true distance to the surface *d*_*s*_(*i*) we calculated the NRMSE in the same way, using 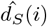 to denote the distance to the surface along the CS-axis:

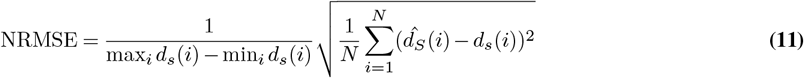

#### Generation of Patterning Maps

To obtain labels that approximate cellular segmentation we used simple linear iterative clustering (SLIC) implemented in scikit-image (63). We restricted the SLIC segmentation to the gastruloid by using a mask derived from the full structure segmentation, and selected a number of superpixels that approximately matched the expected number of cells, based on reported cell diameters in (28).

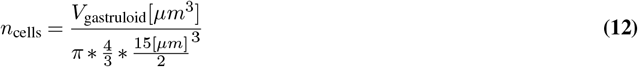

Superpixels were then used to extract intensity-based features from the marker images. For each superpixel, we quantified raw intensities, thresholded signals, marker co-localization, and centroid positions. Thresholds for each marker were manually determined based on multiple representative samples. Co-localization was defined as the overlap of positive voxels in two marker channels. Among these, only co-localization features for T&FOXA2 and FOXA2&SOX17 were used, as they correspond to early mesendoderm or axial mesoderm, and definitive endoderm signatures, respectively. Superpixel centroids were further used to compute normalized anterior-posterior (AP) and cross-sectional (CS) positions, as detailed in Supplemental Note 3. To generate patterning maps, superpixels were grouped into equally spaced bins along the AP and CS axes, and the average thresholded signal or co-localization intensity was calculated for each bin.

#### Uniform Manifold Approximation Projection (UMAP) Dimensionality Reduction and Clustering

To investigate the patterning data, we first reduced its dimensionality using Uniform Manifold Approximation and Projection (UMAP), as implemented in umap-learn (64), using parameters: n_neigbors = 20, min_dist = 0,1, metric = euclidean. For clustering, PCA was performed (see section “Principal Component Analysis”), and the first ten principal components (PCs) of the dataset were used to build a k-nearest neighbor (kNN) graph (FNN, k = 10). Leiden clustering (65) was then conducted as implemented in the leidenbase R package, with a resolution parameter of 0.3. We explored the resulting clusters by interactively inspecting maximum intensity projections of the original input datasets. This exploratory analysis revealed that one of the initial clusters (Cluster 3) encompassed phenotypically distinct subgroups, which also exhibited clear differences in feature space based on principal component analysis (see Supplemental Video 8 and Fig. S3D). To resolve this heterogeneity, cells belonging to the initial Leiden cluster 3 were subsetted, a new kNN graph was constructed from their PCs, and Leiden clustering was repeated with resolution = 0.75. Sub-cluster assignments were then merged back into the full dataset. The resulting 4 clusters were then validated using cluster separation indices computed in PCA space. To ensure robustness, we bootstrapped the dataset 100 times by sampling random subsets comprising 90% of the data. For each iteration, we calculated cluster separation indices including Calinski-Harabasz, Davies-Bouldin, and Silhouette scores. We further combined the scores by ranking each index (with higher rank indicating better separation) and calculated the mean rank per iteration.

#### Principal Component Analysis

Standard scaling of the patterning features was performed prior to fitting principal component analysis (PCA), as implemented in scikit-learn (66). We initially retained the full set of components to examine the explained variance across the dataset. To explore associations between sample-level properties (e.g., volumes and *N*_0_) and principal components, we computed Spearman correlations. In order to interpret the contributions of individual features to each component, we visualized the PCA loadings as patterning heatmaps for each marker.

#### Comparison of 2D maximum projection and SpinePy 3D Patterning Analysis

We first generated synthetic signal fields representing distinct posterior pattern types (for details, see Supplemental Note 1). We then performed superpixel segmentation on the synthetic data to simulate cell-like voxels. These superpixels were used to quantify mean signal intensities within the synthetic fields, and the quantified values were then mapped back to the superpixel labels to yield realistic 3D patterning images (Fig. S2A; Supplemental Videos 4–7). To extract 2D maximum projection based patterning features, we adapted the pipeline described in (28) to be compatible with our data format (original code available at Git-Lab: https://gitlab.pasteur.fr/tglab/gastruloids_precisionandscaling/-/tree/main). Specifically, we used maximum intensity *z*-projections of the gastruloid segmentation to identify the medial axis and define AP axis segments. Signal was then quantified in maximum intensity projections of the remapped synthetic patterns (Fig. S2A, bottom). Extracted profiles were aligned by sorting them from high posterior to low anterior signal and binned into 20 equally spaced AP segments per structure. For 3D patterning quantification, we applied SpinePy to extract the medial axis via skeletonization, and subsequently employed the same AP-based patterning map approach described above. To derive a dimensionality-reduced description of the resulting patterning space, we applied Uniform Manifold Approximation and Projection (UMAP) with n_neighbors = 50 after performing *z*-score normalization. We then assessed the separation of different pattern types using clustering performance metrics implemented in scikit-learn (66), including the Calinski-Harabasz index, Silhouette score, and Davies-Bouldin score. To ensure a fair comparison between methods, we first applied PCA with prior standard scaling to both the 2D and 3D feature spaces, obtaining equal-dimensional representations. Cluster separation was then compared using the known (i.e., the simulated, ground-truth) patterning types as cluster labels and the first 20 principal components as the feature space.

### Statistical Analysis

#### Statistical Assumptions and Tests

Data were first assessed for normality using the Shapiro–Wilk test and for homogeneity of variances using Levene’s test. If data were normally distributed with equal variances and paired, repeated-measures ANOVA was performed using least squares regression, followed by paired t-tests for post hoc comparisons with Benjamini–Hochberg false discovery rate (FDR) correction. For unpaired data under the same assumptions (normal distribution and equal variance), a one-way (between-groups) fixed-effects ANOVA was performed using Type II sums of squares, with Tukey’s Honest Significant Difference (HSD) test for post hoc comparisons. When data violated assumptions of normality or equal variance but were paired, the Friedman chi-squared test was used with post hoc Wilcoxon signed-rank tests, also corrected using the Benjamini–Hochberg procedure. For non-normal or heteroscedastic unpaired data, the Kruskal–Wallis test was applied, followed by Dunn’s test for post hoc comparisons with FDR correction. Categorical data were analyzed using the Chi-squared test, and if significant, post hoc pairwise Chi-squared tests were performed with Benjamini–Hochberg FDR adjustment.

#### Logistic Regression Modeling of Cluster Predictors

To assess the relative contributions of standardized volume and initial cell number (*N*_0_) to the classification of cluster identities, we conducted a series of pairwise logistic regression analyses in R (version 4.1.2). Volumes were standardized (z-scored), and both cluster identities and *N*_0_ were converted to factors. We chose *N*_0_ = 400 — the middle value — as the reference level. For each pairwise comparison between cluster identities, we subset the data accordingly and re-levelled cluster identities such that one of the two clusters served as the reference. Within each pair, we fitted three binomial logistic regression models: i) A model including only standardized volume; ii) A model including only *N*_0_; iii) A full model including both predictors (standardized volume + *N*_0_). Akaike Information Criterion (AIC) values were then extracted to evaluate relative model fit. To assess the statistical contribution of each predictor, likelihood ratio tests (LRTs) were performed: i) between the volume-only model and the full model (testing the added value of *N*_0_); ii) between the *N*_0_-only model and the full model (testing the added value of standardized volume). *P* -values from LRTs were adjusted for multiple testing using the Benjamini–Hochberg procedure. Significance threshold was set at a Holm-adjusted *P* < 0.05.

**Fig. S1.**
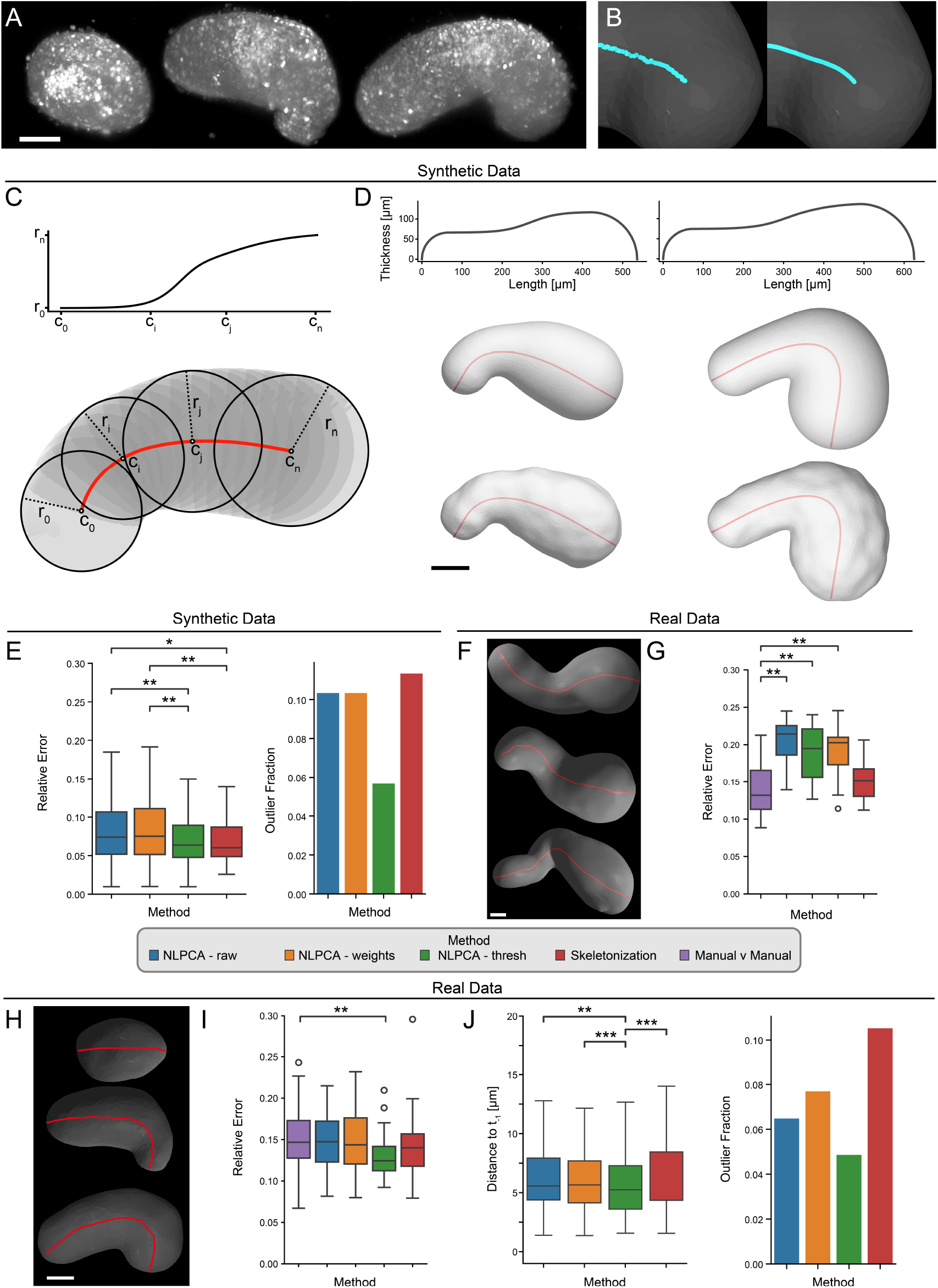
Synthetic and Real Data Spine Extraction Validation: **A** Maximum intensity projections of time-lapse data used to generate data for spine extraction. left: 96 h, middle 103 h, right: 109 h, used for segmentation and surface reconstruction in 1C and D. Scale bar: 100 *µ*m. **B** Visualization of the smoothing procedure. Left: coordinates extracted using 3D skeletonization, right: coordinates after smoothing. Scale bar: 100 *µ*m. **C** Generation of synthetic gastruloids. A random curve is generated (red) with coordinates *c*_0_ , *c*_*i*_, *c*_*j*_ , *c*_*n*_ that have associated radii *r*_0_ , *r*_*i*_, *r*_*j*_ , *r*_*n*_ defined by a sigmoid profile (top) (see Methods). **D** Examples of synthetic gastruloids. Top: radial profiles along the whole axis. Middle: surface reconstructions of synthetic gastruloids generated with the corresponding profiles. Bottom: Reconstructed surfaces with Perlin noise added to surface vertices. Scale bar: 100 *µ*m. **E** Validation of extracted spines with synthetic data. Left: relative error of the extracted spines to the real curves used to generate the data (see Methods) with outliers removed. Right: quantification of the fraction of outliers for each method. Threshold for outliers calculated as 1.5 * 75th percentile using all data. Statistics: Friedman Chi-square *P* < 0.0001. Post-hoc Dunn test with Benjamini-Hochberg FDR correction. **F** Three examples of gastruloids with complex morphologies. AP-axis annotation shown in red. Scale bar: 100 *µ*m. **G** Relative error of detected spines for complex structures. Three experts annotated 11 gastruloids spines which were used to calculate the relative error (see Methods) of the extracted spines to the manual annotations, and of the manual annotations of each user to one another. Statistics: Repeated measures Anova using least squares regression, *P* = 0.000003; post-hoc: two-sided t-test comparing manual-v-manual errors to the spine extraction errors, with FDR-BH correction. **H** Example annotations for a time-lapse dataset. Scale bar: 100 *µ*m. **I** Relative error of detected spines for time-lapse structures. 40 annotations from three experts were used to calculate the normalized distances (see Methods) of the extracted spines to the manual annotations, and of the manual annotations of each user to one another. Statistics: Friedman Chi-square, *P* = 0.0269. Post-hoc: one-sided Wilcoxon tests investigating if spine extraction errors are smaller than manual-v-manual errors, with Benjamini-Hochberg FDR correction. **J** Consistency of extracted spine over time in live imaging data. Left: average distance to the spine path at the previous time point, with outliers removed. Friedman Chi-square *P* = 0.0002. Post-hoc: two-sided Wilcoxon test with Benjamini-Hochberg FDR correction. Right: quantification of the fraction of outliers for each method. Threshold for outliers calculated as 1.5 * 75th percentile using all data. * *P* < 0.05, ** *P* < 0.01 For all boxplots, boxes represent the inter quartile range (IQR), whiskers indicate 1.5 * IQR, line denotes median, circles represent outliers. Sample sizes: **E** *n* = 300; **G** *n* = 11, **I** *n* = 40; **J** *n* = 249.

**Fig. S2.**
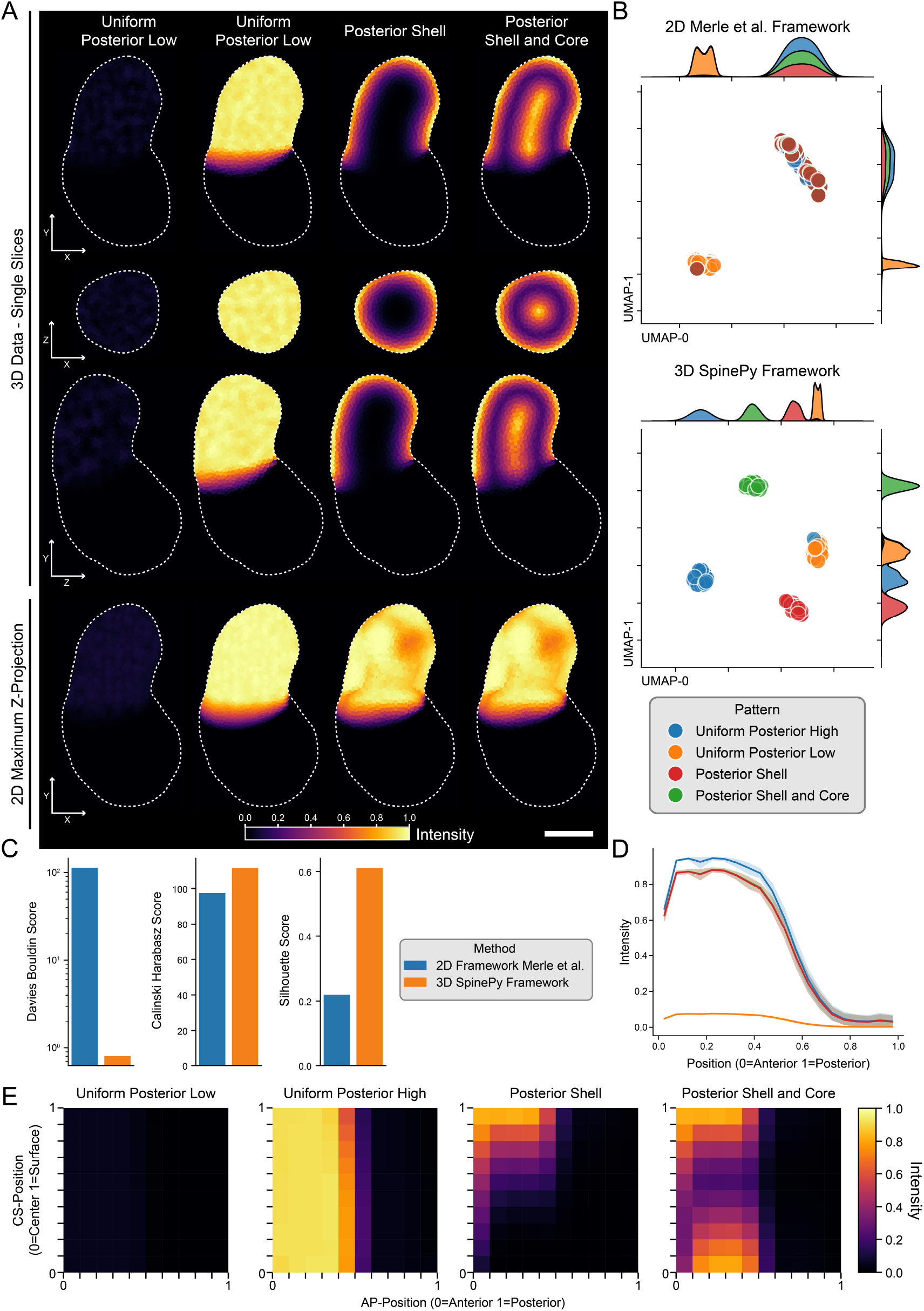
Direct comparison of 2D and 3D Analysis: **A** Representative examples of synthetic patterns. Scale bar: 100 *µ*m. **B** Uniform Manifold Approximation and Projection (UMAP) of profile features extracted using the 2D analysis framework (28) (top) and patterning map features extracted using SpinePy (bottom). Above and to the side density plots, with overlapping density shown as stacked plots. Samples are colored by the simulated patterning class. **C** Cluster separation indices for patterning classes in Principal Component Analysis (PCA) space for 2D and 3D analysis (see Methods). For the Davies Bouldin Score low values indicate better separation. For the Calinsky Harabasz Score and Silhouette Score high values indicate better separation. **D** Average Profile plots extracted using 2D analysis pipeline for each patterning class. Line indicates mean and shaded area 95% confidence interval. **E** Average patterning maps extracted using SpinePy shown for each patterning class. Sample sizes: **B** 2D - *n* = 91; 3D - *n* = 100 **D** *n* = 91 **E** *n* = 100.

**Fig. S3.**
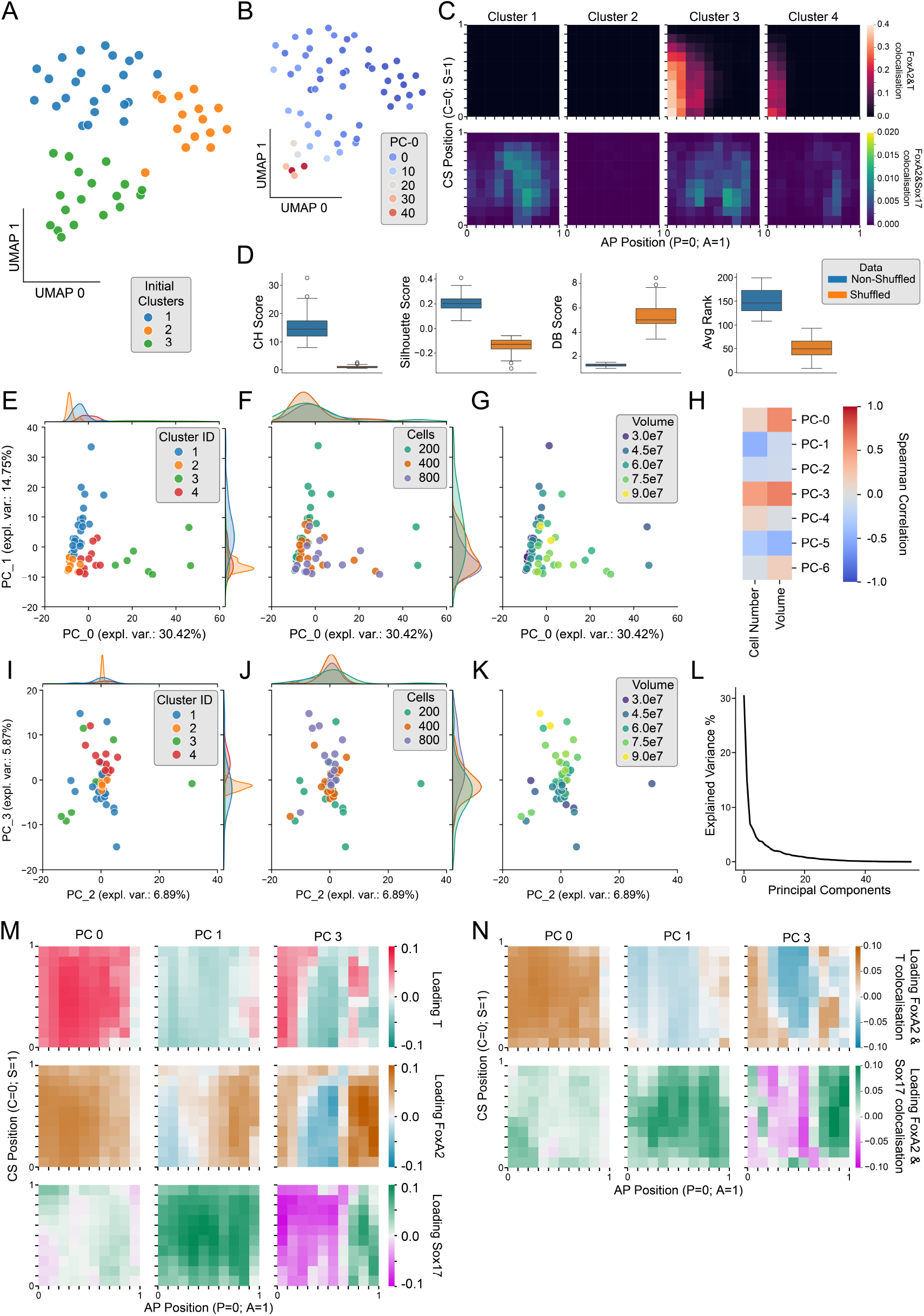
UMAP and PCA analysis of patterning maps: **A** Uniform Manifold Approximation and Projection (UMAP) of patterning map features overlaid with *k*-means clustering results with *k* = 3. **B** UMAP of patterning map features with PC-0 (determined using Principal Component Analysis (PCA)) values overlaid. **C** Box plot showing cluster separation scores for non-shuffled versus shuffled data across 100 bootstrap iterations. For each iteration, Calinski-Harabasz (CH), Davies-Bouldin (DB), and Silhouette scores were computed in PCA space. To summarize performance across indices, we ranked clusters per metric (higher rank = better separation) and computed the mean rank (Avg Rank) per iteration. **D** Average patterning maps for the co-localization of indicated developmental markers, separated per cluster. **E** Middle: PC-0 and PC-1, colored by clusters obtained after sub-clustering initial Cluster 3. Top & right: density plots for PC-0 and PC-1, respectively. **F** Middle: PC-0 and PC-1, colored by initial cell number (*N*_0_ ). Top, right: density plots for PC-0 and PC-1, respectively. **G** PC-0 and PC-1, colored by structure volume at the culture end-point (*t* = 120*h*). **H** Heatmaps showing Spearman correlations of indicated principle components with initial cell number (*N*_0_ ) and total end-point gastruloid volume. **I** Middle: PC-2 and PC-3 with clusters obtained after sub-clustering initial Cluster 3. Top, right: density plots for PC-2 and PC-3, respectively. **J** Middle: PC-2 and PC-3 , colored by initial cell number (*N*_0_ ). Top, right: density plots for PC-2 and PC-3, respectively. **K** PC-2 and PC-3, colored by structure volume at the culture end-point (*t* = 120*h*). **L** Explained variance percentage for all principal components. **M**,**N** PCA loadings visualized as patterning maps for developmental markers **L** and co-localization of developmental markers, as indicated **M**. Sample sizes: **A** Cluster 1 - *n* = 23, Cluster 2 - *n* = 14, Cluster 3 - *n* = 19; **B D, H** Cluster 1 - *n* = 23, Cluster 2 - *n* = 14, Cluster 3 - *n* = 6, Cluster 4 - *n* = 13; **E, I** *N*_0_ = 200 - *n* = 20, *N*_0_ = 400 - *n* = 20, *N*_0_ = 800 - *n* = 16.

