## Supplemental Video Captions for "SpinePy enables automated 3D spatiotemporal quantification of multicellular *in vitro* systems"

### Supplemental Video Captions to "SpinePy enables automated 3D spatiotemporal quantification of multicellular *in vitro* systems"

**Ryan G. Savill**[1,2,✉], **Alba Villaronga-Luque**[1,2,‡], **Marc Trani Bustos**[1,2,3,‡], **Yonit Maroudas-Sacks**[1,3], **Julia Batki**[4], **Alexander Meissner**[4,5], **Allyson Q. Ryan**[1,6,§], **Carl D. Modes**[1,3,6], **Otger Campàs**[1,3], and **Jesse V Veenvliet**[1,3,6,✉]

[1]Max Planck Institute of Molecular Cell Biology and Genetics, Dresden, Germany
[2]Faculty of Biology, Technische Universität Dresden, Dresden, Germany
[3]Cluster of Excellence Physics of Life, Technische Universität Dresden, Dresden, Germany
[4]Max Planck Institute for Molecular Genetics, Berlin, Germany
[5]Department of Biology, Chemistry and Pharmacy, Freie Universität Berlin, Berlin, Germany
[6]Center for Systems Biology Dresden, Dresden, Germany
[§]Present address: Chan Zuckerberg Biohub, 499 Illinois Street, San Francisco, CA 94158
[‡]These authors contributed equally to this work and should be considered shared second author

#### Supplemental Video 1 – spine extraction comparison for 3D live imaging of gastruloids

All extracted spine paths are shown in cyan and surface meshes are displayed in a translucent gray. **top left:** skeletonization approach. Image skeleton is shown in orange **top right:** non-linear principle component analysis (NLPCA) approach using all seed points (displayed as orange points). **bottom left:** NLPCA approach using weighted seed points (displayed with a viridis colormap, purple - close to surface, yellow - close to center). **bottom right** NLPCA approach using a threshold to filter out points closer to the surface (seed points shown in orange. Scale bar $100\,\mu m$

#### Supplemental Video 2 – density gradient measurement in synthetic data

Surface mesh shown in gray, spine path shown in red, density field shown in purple, and synthetic density points shown in green. The sliced mesh is shown for every section along the AP axis to demonstrate which points are used to calculate the density of a section. Scale bar $50\,\mu m$

#### Supplemental Video 3 – determination and quantification of normalized spatial information

Surface mesh shown in gray, spine path shown in red, point of interest shown in yellow. Anteroposterior (AP location indicated in red and core-to-surface (CS) position in shown in blue. Superpixel segmentation is overlaid on nuclear signal (DAPI) before normalized AP position and CS positions are shown by coloring superpixels by their location (dark - posterior / center - bright anterior / surface). Lastly the quantification of TBXT is shown in 2D and 3D. Scale bar $100\,\mu m$

#### Supplemental Videos 4 - 7 – synthetic patterning examples

Synthetic generated patterns shown by moving through the z - planes. Gastruloid boundary shown in white and synthetic signal shown with the inferno colormap (purple - low signal, yellow - high signal). Top left: x/y plane, top right: z/x plane, bottom left z/y plane. Scale bar $100\,\mu m$. **4:** low intensity pattern, **5:** high intensity pattern, **6:** shell pattern, **7:** shell and core pattern

#### Supplemental Video 8 – interactive investigation of initial unsupervised clustering

UMAP of the patterning map feature space colored by initial clusters (blue: cluster 1, orange: cluster 2, green: cluster 3). When hovering over datapoints the corresponding maximum z-projections of the input data is shown where the stainings are visualized: nuclei (gray DAPI), TBXT (red), FoxA2 (yellow) and SOX17 (blue).
