## Supplemental Notes for "SpinePy enables automated 3D spatiotemporal quantification of multicellular *in vitro* systems"

### Supplemental Note 1 Synthetic Data Generation

**Synthetic Spine Paths.** To simulate gastruloids we start by generating curves in 3D space with varying degrees of curvature and randomness as a baseline to simulate the AP axis:

$$\mathbf{c}(s) = \begin{pmatrix} x(s) \\ y(s) \\ z(s) \end{pmatrix} = s \begin{pmatrix} \cos\big(o_x + \kappa a_x s\pi\big) \\ \sin\big(o_y + \kappa a_y s\pi\big) \\ \sin\big(o_z + \kappa a_z s\pi\big) \end{pmatrix} \mathbf{S}_{\text{img}} \quad s \in [0,1]; \mathbf{o} \qquad \sim \mathcal{U}\big([0,2\pi]^3\big), \mathbf{a} \qquad \sim \mathcal{U}\big([0.5,1.5]^3\big). \tag{1}$$

With $\mathbf{o}$ and $\mathbf{a}$ being an offset vector and a curvature vector, respectively, both drawn from a random uniform distribution $\mathcal{U}$ and $\kappa$ being a global curvature factor that determines the degree of curvature. The scaling and centering matrix $\mathbf{S}_{\text{img}}$ then transforms and scales the path to be centered in an image with size (500, 500, 500) and scaled until it reaches bounds defined by bounding box argument. We generated 3 datasets with less curved gastruloids $\kappa = 0.08$, more curved gastruloids $\kappa = 0.3$ and a dataset that lies in the z-plane. The more curved gastruloids had an additional curvature multiplier $\mathbf{k}$ added:

$$\mathbf{c}_{curved}(s) = \begin{pmatrix} x(s) \\ y(s) \\ z(s) \end{pmatrix} = l\, s \begin{pmatrix} \cos\big(o_x + \kappa k_x a_x s\pi\big) \\ \sin\big(o_y + \kappa k_y a_y s\pi\big) \\ \sin\big(o_z + \kappa k_z a_z s\pi\big) \end{pmatrix} \mathbf{S}_{\text{img}} \quad s \in [0,1]; \tag{2}$$

With $\mathbf{k}$ being a vector of 3 factors that are drawn from $[1,1.5]$ with a probability of 0.5. For the Dataset with curves lying in the z-plane we simply define the curves as:

$$\mathbf{c}(s) = \begin{pmatrix} x(s) \\ y(s) \\ z(s) \end{pmatrix} = l\, s \begin{pmatrix} \cos\big(o_x + \kappa a_x s\pi\big) \\ \sin\big(o_y + \kappa a_y s\pi\big) \\ 0 \end{pmatrix} \mathbf{S}_{\text{img}} \quad s \in [0,1]; \tag{3}$$

After generating curves we sample $n$ points along the curve so that each point $c_i$ is spaced 1 unit distance from each other:

$$\mathbf{c}_i = \mathbf{c}\big(\tfrac{i}{n}\big), \quad i = 0,\ldots,n \tag{4}$$

**Synthetic Thickness Profiles.** To synthesize gastruloids the next step is generating a thickness profile along the curve by defining random start and end radii and blending between them with a sigmoid profile:

$$r_i = r_{\text{start}} + \big(r_{\text{end}} - r_{\text{start}}\big)\,\text{sigmoid}(i), \quad r_{\text{start}} \sim \mathcal{U}\big(57,95\big), \quad r_{\text{end}} \sim \mathcal{U}\big(88,147\big) \tag{5}$$

We opted to use a sigmoid profile centered at a midpoint $m$ drawn from a uniform distribution with some adjustments:

$$\text{sigmoid}(i) = \frac{1}{1+\exp\big(-\alpha(i)\,(\frac{i}{n} - m)\big)}, \quad m \sim \mathcal{N}(0.5, 0.1) \tag{6}$$

The steepness factor $\alpha$ is a function that transitions between two steepness factors $\alpha_1$ and $\alpha_2$ (also drawn from uniform distributions), blending their values around the midpoint to avoid kinks:

$$\alpha(i) = \alpha_1\big(1-\sigma_{20}(i)\big) + \alpha_2\sigma_{20}(i),\ \sigma_{20}(i) = \frac{1}{1+\exp\big(-20(\frac{i}{n} - m)\big)}, \quad (\alpha_1,\alpha_2) \sim \mathcal{U}\big(5,15\big) \tag{7}$$

Each point $\mathbf{c}_i$ on the curve now has a profile radius $r_i$ associated to it.

**Synthetic Surface Mesh Generation.** We use the profile to generate a distance field $\mathrm{D}$ in the 3D image (size 500,500,500) as:

$$\mathbf{x} = (x,y,z)^\top,\ \mathrm{D}(\mathbf{x}) = \min_{0\le i\le n} \frac{\|\mathbf{x} - \mathbf{c}_i\|}{r_i} \tag{8}$$

We can then obtain a label image through binarization of the distance field into a label image $\mathrm{L}(\mathbf{x}) = \mathrm{D}(\mathbf{x}) < 1$. Following binarization we reconstruct the surface analogously to the time-lapse data. Since these surfaces are very smooth we add some Perlin noise to the vertices to mimic real data. First large scale noise is added to the surface vertex normals (with a factor between -7.5 and 7.5) to simulate large scale fluctuations in width before adding small scale noise (with a factor between -3.5 and 3.5) to simulate cell-scale noise.

**Synthetic Density Gradients.** To validate the scalar field quantification we investigated density profiles in synthetic gastruloids. To generate uniform density gradients we randomly sample points within the bounding box of the gastruloid surface and sequentially include points lying within the surface mesh until a specified total density level $\rho_{syn}$ is reached. We chose densities of $10^{-3}$, $10^{-4}$ and $10^{-5}$ to cover a range of expected density levels.
To quantify gradient density profiles we used a dateset of synthetic gastruloids, for which the axis does not change it's z-position. This allowed us to create a location map in 2D which defined 10 location segments along the AP-axis, indicated by the pixel values $\mathrm{I}(x,y)$, $\mathrm{I}(x,y) \in [0,1]$, $(0 = posterior, 1 = anterior)$. This location map was blurred with a gaussian blur filter (sigma = 20) and could be used to generate density maps P spanning density values between $\rho_{start}$ and $\rho_{end}$:

$$\mathrm{P}(x,y) = \rho_{\text{start}} + I(x,y)\big(\rho_{\text{end}} - \rho_{\text{start}}\big) \tag{9}$$

**Synthetic Patterns.** For synthetic patterns we decided to emulate posterior marker gene expression. To generate these realistic synthetic patterns we first required positional map A to define the anterior-posterior location. Similarly to the distance map we can obtain this map by utilizing the synthetic curves and radial profiles, defined in equations 1 and 5:

$$\mathrm{A}(\mathbf{x}) = \frac{1}{n} \operatorname*{argmin}_{0 \le i \le n} \frac{\|\mathbf{x} - \mathbf{c}_i\|}{r_i}, \quad \mathbf{c}_i = \mathbf{c}\big(\tfrac{i}{n}\big), \quad i = 0, \dots, n \tag{10}$$

This map will assign every position in 3D space to an anterior-posterior location. We then mark the posterior region with another map T, which indicates posterior positions with 1 and non-posterior positions with 0. We can then set a threshold $\tau = 0.45$ for which parts of the posterior we want to use to generate patterns and also add a crossover region in which the posterior signal fades by defining:

$$\mathrm{T}(\mathbf{x}) = \max\Big(u(\tau - \mathrm{A}(\mathbf{x})),\ \mathrm{clip}(-5 \cdot \|\mathrm{A}(\mathbf{x}) - \tau\|) + 1\Big), \quad u(z) = \begin{cases} 1, & z > 0, \\ 0, & z \le 0. \end{cases} \tag{11}$$

with $\mathrm{clip}$ representing a clipping function that clamps values between 0 and 1 defined as:

$$\mathrm{clip}(x;0,1) = \begin{cases} 0, & x \le 0, \\ x, & 0 < x < 1, \\ 1, & x \ge 1. \end{cases} \tag{12}$$

The first term in the $\max$ operator represents the posterior region defined by $\tau$, whereas the second term is a gradient peaking at 1 for positions close to $\mathbf{c}(\tau)$ and fading sharply as the distance increases. This gives us a posterior region with a small transitional zone. We can now use this map to define uniform low ($\mathrm{I}_{low}$) and high ($\mathrm{I}_{high}$) intensity posterior regions as:

$$\mathrm{I}_{low}(\mathbf{x}) = \mathrm{T}(\mathbf{x})\big(0.05 + \mathcal{P}(\mathbf{x})\big), \quad \mathrm{I}_{high}(\mathbf{x}) = \mathrm{T}(\mathbf{x})\big(0.95 + \mathcal{P}(\mathbf{x})\big), \tag{13}$$

with $\mathcal{P}(\mathbf{x}) \in [-0.05, 0.05]$ being a Perlin noise function that returns random values that are smooth in local space.
To generate a posterior shell pattern with high intensity at the surface and low intensity inside we use the distance map $D$ that we defined in Equation 8. Since values inside the gastruloid are all below 1 we can make a steep gradient $G$ from the outside to the inside, constrained to the posterior region by defining:

$$\mathrm{G}(\mathbf{x}) = \mathrm{clip}\big(\mathrm{D}(\mathbf{x})^3\big)\mathrm{T}(\mathbf{x}) \tag{14}$$

with $\mathrm{clip}$ representing the clipping function defined in equation 12. Lastly the shell and core pattern is generated by combining a core rod-like pattern as:

$$\mathrm{C}(\mathbf{x}) = \mathrm{clip}(-\mathrm{D}(\mathbf{x}) + 1)^2\mathrm{T}(\mathbf{x}) \tag{15}$$

and taking the maximum value of the shell (G) or the core pattern (C).

### Supplemental Note 2: Slice Optimization

To determine the initial planes for slicing we decide on a number of volume sections $n_{\text{sec}}$ and then determine $n_{\text{planes}} = n_{\text{sec}} - 1$ equally spaced points $\mathbf{p}_j$ along the spine (excluding the poles):

$$\mathbf{p}_j = \mathbf{c}(s_j),\; s_j = \frac{j}{n_{\text{sec}}} \tag{16}$$

and their corresponding tangents, which form the plane normals $\mathbf{n}_j$ as:

$$\mathbf{n}_j = \frac{\mathbf{c}'(s_j)}{\|\mathbf{c}'(s_j)\|} \tag{17}$$

To calculate the loss for intersecting slices for each plane $j$ we first cast $n_{\text{ray}}$ rays in a circle along the plane defined as:

$$\mathcal{R}_{j,i} = \left\{\mathbf{p}_j + t\,\mathbf{R}\big(\mathbf{n}_j, \phi_i\big)\mathbf{u}_j \;\middle|\; t \geq 0\right\},\; \phi_i = \tfrac{2\pi i}{n_{\text{ray}}} \tag{18}$$

with $\mathbf{u}_j \cdot \mathbf{n}_j = 0, \|\mathbf{u}_j\| = 1$. For each ray we record the first intersection with the surface mesh $\mathbf{q}_S$ to calculate the distance to the surface along the ray:

$$d_S = \|\mathbf{p}_j - \mathbf{q}_S(\mathcal{R}_{j,i})\| \tag{19}$$

We then measure the first intersections with other planes $\mathbf{q}_m$ parametrized by $\mathbf{p}_m$ and $\mathbf{n}_m$, $m = 0, \ldots, n_{\text{planes}}$, $m \neq j$ to determine the distance to the intersection as:

$$d_I = \begin{cases} d_S, & \text{if } \mathbf{q}_m(\mathcal{R}_{j,i}) \text{is outside the surface} \\ \|\mathbf{p}_j - \mathbf{q}_m(\mathcal{R}_{j,i})\|, & \text{otherwise} \end{cases} \tag{20}$$

making sure that all intersection distances outside of the surface are not taken into account. The loss $L$ for each ray is then calculated as:

$$L_{j \to m}(i) = \frac{d_S - d_I}{d_S} \tag{21}$$

The final loss for all slices $\mathcal{L}$ is the sum of all individual ray losses $L$, as well as an additional loss term penalizing rotation of the plane normals, so that optimized slices are still close to orthogonal with the spine:

$$\mathcal{L} = \sum_{j \neq m} \sum_i L_{j \to m}(i) \mathcal{L} = \sum_{j \neq m} \sum_i L_{j \to m}(i) + \lambda\, \mathbf{n}_j \cdot \mathbf{n}_j^{(0)} \tag{22}$$

with $\mathbf{n}_j^{(0)}$ representing the un-rotated plane normal corresponding to the tangent of the curve at $\mathbf{c}_j$. This additional factor $\lambda$ allows us to penalize plane rotations. This is crucial to keep planes as close to orthogonality as possible, since without this factor a valid optimization solution will be to set all planes parallel to each other.

### Supplemental Note 3: Location Measurements to Determine a Spine-Based Coordinate System

Since we want to define a common coordinate system for all gastruloids we have to be able to map ROIs from 3D space into a common space. An obvious choice would be a cylindrical coordinate system in which we can describe the location of region of interest (ROI) as a function of three parameters:

$$\mathbf{l}(s,r,\phi) = \mathbf{c}(s) \, + \, r\,R(s)\big[\cos\phi\,\mathbf{n}(s) + \sin\phi\,\mathbf{b}(s)\big], \quad 0 \le s \le 1,\ 0 \le r \le 1,\ \phi \in [0,2\pi). \tag{23}$$

which would correspond to the AP-axis position ($\mathbf{c}(s)$), radial-position $r$ (from here on core-to-surface position, CS-axis) as a fraction of the distance to the surface $R(s)$, and the rotational-position (represented by $\phi$). $\mathbf{n}(s)$ and $\mathbf{b}(s)$ are two orthogonal vectors that are also orthogonal to the tangent$\mathbf{t}$ of the spine at position $s$, that allow us to draw the vector to the surface, defined as:

$$\mathbf{t}(s) = \frac{\mathbf{c}'(s)}{\|\mathbf{c}'(s)\|},\ \mathbf{n}(s) \perp \mathbf{t}(s),\ \mathbf{b}(s) = \mathbf{t}(s) \times \mathbf{n}(s). \tag{24}$$

In lack of a common reference position to construct $\mathbf{b}$ and $\mathbf{n}$ along the entire AP-axis we further simplify this coordinate system by treating gastruloids as rotationally symmetric, thus reducing the coordinate system to:

$$\mathbf{l}(s,r) = \mathbf{c}(s) \, + \, r\,R(s) \tag{25}$$

For each ROI position $\mathbf{l}$ we thus only need to find the corresponding axis coordinate $\mathbf{c}(s)$ and construct the vector between $\mathbf{c}(s)$ and the ROI to determine $R(s)$. To detect a suitable value for $s$ for a ROI we have two available approaches. The first determines the closest point along the AP axis numerically, by quantifying all distances from the ROI to all axis points $\mathbf{c}(s),\ s \in [0,1]$ as:

$$d_{\text{AP}}(s) = \|\mathbf{c}(s) - \mathbf{l}\| \tag{26}$$

and then determining the optimal value for s by minimization:$10^{-3}$:

$$s^* = \underset{s\in[0,1]}{\operatorname{argmin}}\, d_{\text{AP}}. \tag{27}$$

Since surface intersections are computationally cheap when employing optimized ray casting we solve this minimization numerically with a resolution of 0.0001:

$$\Delta s = 10^{-3},\ \mathcal{S} = \{k\Delta s \mid k = 0,\ldots,10^3\},\ \tilde{s} = \underset{s\in\mathcal{S}}{\operatorname{argmin}}\, d_{\text{AP}}. \tag{28}$$

The second approach minimizes not only the distance to the spine $d_{AP}(s)$ but also the distance to the surface intersection $\mathbf{q}(s)$ along the vector between the spine and ROI $d_S(s)$:

$$d_{\text{comb}}(s) = d_{\text{AP}}(s) + d_{\text{S}}(s), \quad d_{\text{S}}(s) = \|\mathbf{q}(s) - \mathbf{l}\| \tag{29}$$

Using this combined distance we again determine the optimal $s$ through minimization:

$$s^* = \underset{s\in[0,1]}{\operatorname{argmin}}\, d_{\text{comb}}(s). \tag{30}$$

and compute the solution numerically as:

$$\Delta s = 10^{-3},\ \mathcal{S} = \{k\Delta s \mid k = 0,\ldots,10^3\},\ \tilde{s} = \underset{s\in\mathcal{S}}{\operatorname{argmin}}\, d_{\text{comb}}(s). \tag{31}$$

The CS-distance $r$ can then be calculated with:

$$r = \frac{d_{\text{AP}}(s)}{R(s)},\ R(s) = \|\mathbf{c}(s) - \mathbf{q}(s)\| \tag{32}$$
